# Optimal treatment for drug-induced cancer persisters involves release periods and intermediate drug doses

**DOI:** 10.1101/2024.11.29.626082

**Authors:** Mattia Corigliano, Arianna Di Bernardo, Marco Cosentino Lagomarsino, Simone Pompei

## Abstract

Targeted cancer therapies often induce a reversible drug-tolerant state in subpopulations of cells, akin to bacterial persistence. Precise characterization of these “cancer persisters” may inform the design of more effective treatment strategies. A previous investigation into the transition to persistence of colorectal cancer cell lines has revealed a distinct dependence on drug presence and concentration, not typical of bacterial systems. Leveraging these findings, this study uses mathematical modeling to explore intermittent treatment protocols aimed at diminishing the long-term fitness of the treated population, thereby enhancing therapeutic efficacy. We adapt a mathematical model originally designed for bacteria to describe colorectal cancer population dynamics in response to a series of treatment and release cycles. The model predicts the long-term increase or reduction of a treated population, as well as its asymptotic composition, leading us to identify success and failure regions within a clinically accessible parameter space, also in combination with hypothetical drugs that act on persisters. Strikingly, our analysis suggests, perhaps counter-intuitively, that optimal treatment outcomes may be achieved in correspondence of non-zero recovery periods and lower than currently administered in the clinics drug concentrations. Furthermore, by incorporating patient drug pharmacokinetics in the model, we demonstrate that intermittent dosing strategies currently explored in clinical trials can be optimized to potentially rival the efficacy of continuous dosing regimens. These findings underscore the potential of mathematical models in guiding the design of optimal treatment protocols by fine-tuning non-trivial decisional trade-offs.

## I. INTRODUCTION

Persistence refers to the ability of a sub-population of cells to survive exposure to a fully inhibitory drug concentration. In bacteria, it has been observed since the early 1940s [1], when researchers first noted that a small fraction of bacterial cells could survive antibiotic treatment while the majority were eliminated. Interestingly, populations grown from these surviving cells exhibited the same behavior, ruling out genetic resistance as the underlying cause. Instead, persistence involves a transient and reversible phenotypic state in which a subset of cells enters a dormant or slow-growing phase, enabling them to evade the lethal effects of antibiotics.

The presence of persister cells poses significant challenges in treating bacterial infections. Their resilience can complicate or undermine treatment, resulting in chronic, recurrent infections, prolonged illness, and the need for more aggressive treatment strategies—when such strategies are available [2]. Crucially, the outcome of a treatment in presence of such phenotypic switches depends sensitively on quantitative parameters of the population dynamics, such as growth, susceptibility and transition rates. Hence, persistence and similar phenotypic switches have inspired extensive mathematical modeling efforts to identify the conditions that contribute to the emergence of resilient subpopulations [2–6].

In recent years, cancer research — leveraging work on cell lines and organoids, and parallels with patient outcomes — has uncovered numerous instances of persistence, revealing a striking similarity between the survival strategies of bacteria and residual tumor cells. Both systems harbor subpopulations of slow-growing persister cells exhibiting remarkable resilience to a wide range of therapeutic agents [7–10]. For example, exposure to certain targeted therapies can reproducibly induce the emergence of *bona fide* persister cells within tumor populations [7, 11, 12]. Crucially, like bacterial persisters, these tumor cells are not genetically resistant to treatment. Instead, they rely on adaptive survival mechanisms that enable them to withstand otherwise lethal therapies. The resurgence of persister cells can drive tumor regrowth, mirroring the case of treated bacteria. This phenomenon has been observed across various tumor types [8], suggesting it may be a general feature, although specific properties may vary between systems. Moreover, persistence in tumors often coexists with more complex, multicellular strategies that enhance therapy evasion.

However, under certain conditions and in specific experimental systems, cancer persisters can be isolated and accurately quantified [12]. At a quantitative level, both bacterial and cancer persisters exhibit a similar biphasic survival pattern, characterized by an initial rapid decline in susceptible cell numbers, followed by a much slower decline corresponding to the surviving persister population (Fig. 1A). Mathematical modeling techniques originally developed to study bacterial persistence have proven adaptable for interpreting the survival curves of cancer cells [12, 13]. Building on this, we previously quantified the transition-to-persistence dynamics in two colorectal cancer (CRC) cell lines (named WiDr and DiFi) when treated with clinical doses of cetuximab, a monoclonal antibody that inhibits EGFR, used in patients with metastatic CRC tumors lacking RAS and BRAF mutations [12]. Cetuximab was administered either alone (in two DiFi cloned populations, called cl. B3 and cl. B6) or in combination with the BRAF inhibitor Dabrafenib (in WiDr cloned populations cl. B5 and cl. B7), used for CRC harbouring BRAF mutations (see appendix A).

**FIG. 1.**
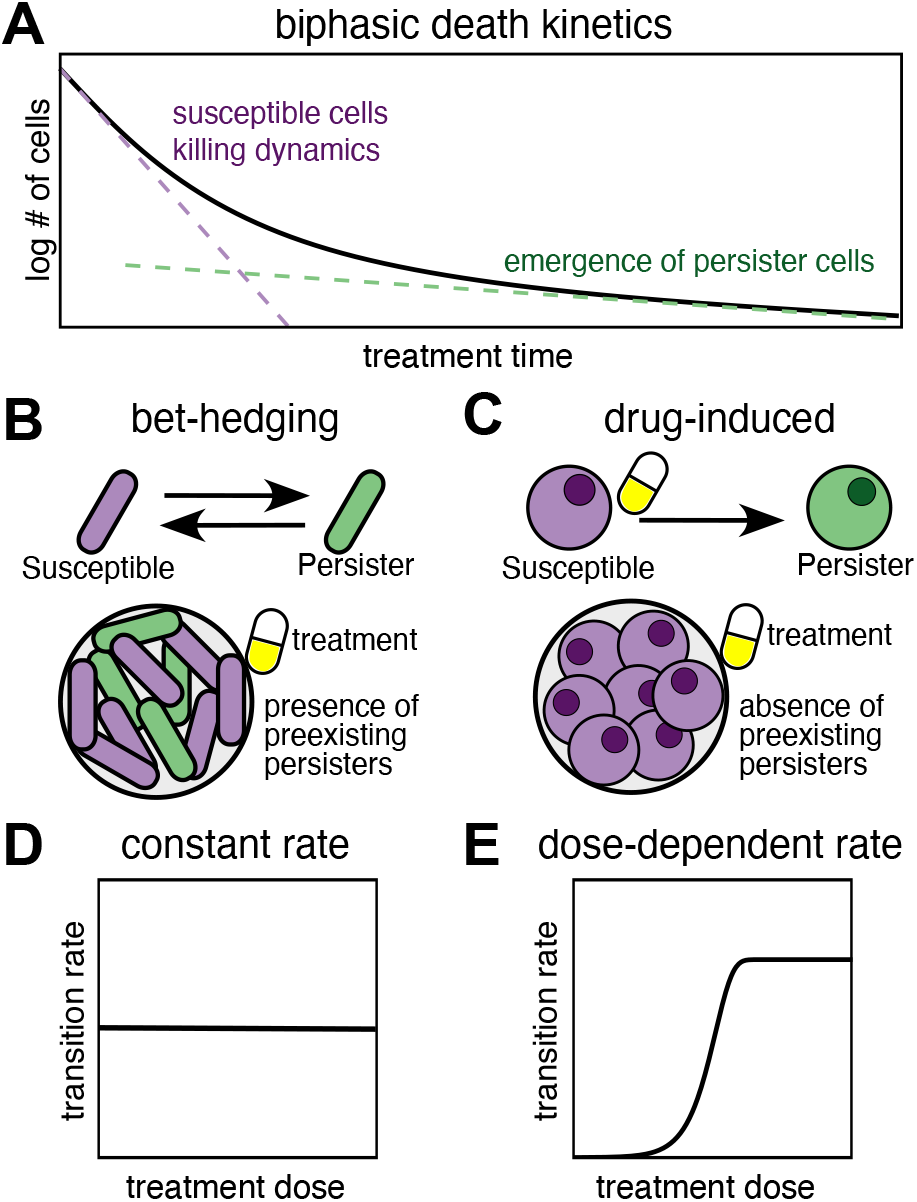
Colorectal cancer persister cells show different dynamics from typical bacterial persisters. **(A)** Biphasic survival curve observed in treated bacterial and cancer cells, characterized by an initial rapid decline in cell numbers followed by a slower phase. This two-timescale death kinetics distinctly indicates the emergence of drug-tolerant persister cells during treatment. **(B)** In the bet-hedging scenario, the transition to persistence occurs independently of the external stress. As a consequence persister cells are already present at the beginning of treatment. **(C)** In the drug-induced persistence scenario, no persisters are expected to be present at the start of treatment, with persister cells emerging only after drug exposure. **(D)** A constant transition rate as a function of drug concentration, typical in cases with weak dependence on drug levels.**(E)** Expected curve of transition rate versus drug concentration observed in colorectal cancer cell lines in ref. [12], showing a monotonic increase that saturates beyond a critical concentration, capturing the drug dependence of persistence in these systems.

Our results demonstrated that in these systems, the persister phenotype was largely induced by drug exposure, rather than pre-existing within the tumor population [12]. By contrast, in bacteria, the transition to a persister state typically occurs spontaneously, reflecting a bet-hedging strategy in face of fast-changing environments (Fig. 1B-C) [5, 9, 14, 15]. Conversely, cancer cells are not evolved to respond to chaging environments, and presumably persistence is related to entirely different physiological behaviors [8, 16]. Additionally, CRC cell-line data sometimes supported a model where the transition rate from normal to persister cells is dependent on drug concentration (Fig. 1D-E). Specifically, the WiDr cell lines, showed a linear increase in the transition rate with rising doses of dabrafenib. Conversely, in the DiFi cell lines, the transition rate was zero in the absence of cetuximab but remained constant across the range of all tested concentrations for treated cells. Notably, the transition rates in both cases can be described by the same mathematical equation: a monotonically increasing function that starts from zero and saturates to a constant value above a critical concentration (Fig. 1E). This critical value is lower than the cetuximab treatment concentrations for the DiFi cell line, while it lies within the range of the investigated dabrafenib concentrations for the WiDr cell line. While these behaviors are intriguing, we currently lack any insight on their long-term impact on treatment.

To address this issue, this study investigates how the quantified features of CRC persisters, particularly their drug-induced nature and the dependence of transition rates on drug concentration, can be leveraged to design optimal drug delivery protocols. This is explored in the context of cyclic treatments, where a series of treatment-release cycles are applied, closely mirroring the therapeutic regimens commonly used in clinical practice [17– 19]. Previous work allows us to direct and restrict our analysis to population dynamics regimes and parameters within the observed range of behaviors [12]. Our modeling framework builds upon previous modeling studies in the context of bacterial persisters [4, 5, 15]. These studies also connect experimental observations to dynamic parameters, giving an insight even when detailed mechanisms remain unclear. Recent work has extended these approaches to a cancer context, though different from the case of persisters we address here [20], highlighting the potential of these tools in guiding clinical treatment strategies. This framework raises several key considerations regarding the design of optimal treatment protocols. For instance, how does the sensitivity of transition rates to drug concentration influence treatment outcomes? Specifically, when transition rates exhibit a weak dependence on drug concentration, it might be possible to optimize the duration of drug release. Alternatively, when transition rates show a strong dependence, one might identify an optimal drug dose to minimize the long-term fitness of treated cancer cells. Finally, one would wish to apply these insights to clinically relevant scenarios, incorporating within-patient, time-dependent drug profile data, to determine whether cyclic treatment protocols offer advantages over constant dosing strategies.

## II. RESULTS

### A. A phenotypic-switch model quantifies the long-term efficacy of cancer-cell periodic treatments

The model describes the dynamics of cancer cell populations under intermittent treatment protocols [3– 5, 15, 20]. An intermittent treatment protocol is characterized by the number of treatment-release cycles, *q*, their respective durations, *τ*_*t*_ and *τ*_*r*_, and the administered dose profile during treatment, *c*(*t*) (Fig. 2A). In this work, we consider protocols that either vary the treatment and release durations, while keeping the dosage constant at clinically-relevant levels, or adjust the dosage and the release time, while keeping constant the treatment duration. Since we aim to quantify and compare the efficacy of all these different treatment protocols, our analysis focuses on the long-term dynamics of the treated cancer population (Fig. 2B), which, as shown in previous work [4, 5], can be used as a valid metric for quantifying protocols’ efficacy.

**FIG. 2.**
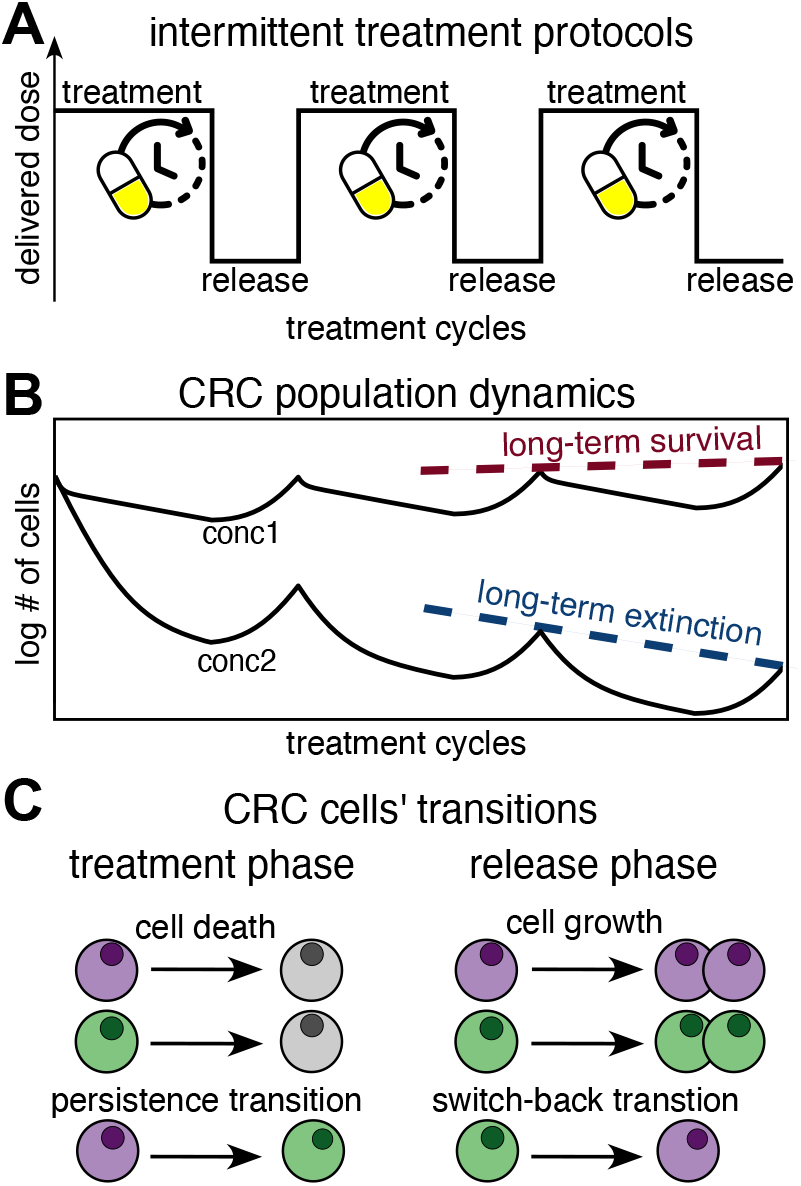
A phenotypic-switch model quantifies the long-term efficacy of periodic treatments. **(A)** A periodic treatment protocol is defined by (i) the number of treatment-release cycles *q*, (ii) the dose profile *c*(*t*) during treatment (constant in the figure), and (iii) the durations of the treatment and release phases, *τ*_*t*_ and *τ*_*r*_. **(B)** Illustration of the predicted cell population dynamics over *q* treatment-release cycles. For fixed durations of treatment and release, the model generates distinct trajectories for different drug concentrations. The long-term behavior, indicated by the asymptotic trend for large *q* (dashed lines), constitutes a proxy for the efficacy of various protocols across many cycles. **(C)** Schematic representation of the key cellular transitions considered in the model. Susceptible (purple) and persister (green) cells can either proliferate or die (rates *μ*_*s*_ and *μ*_*p*_), and transition to the other phenotype (rates *λ*_*s*_, *λ*_*p*_, respectively). These parameters differ between the treatment (superscript *t*) and release (superscript *r*) phases, resulting in a total of eight distinct parameters. Numerical values of the model parameters inferred in [12] and used in our analysis are reported in Table I.

**TABLE I.**
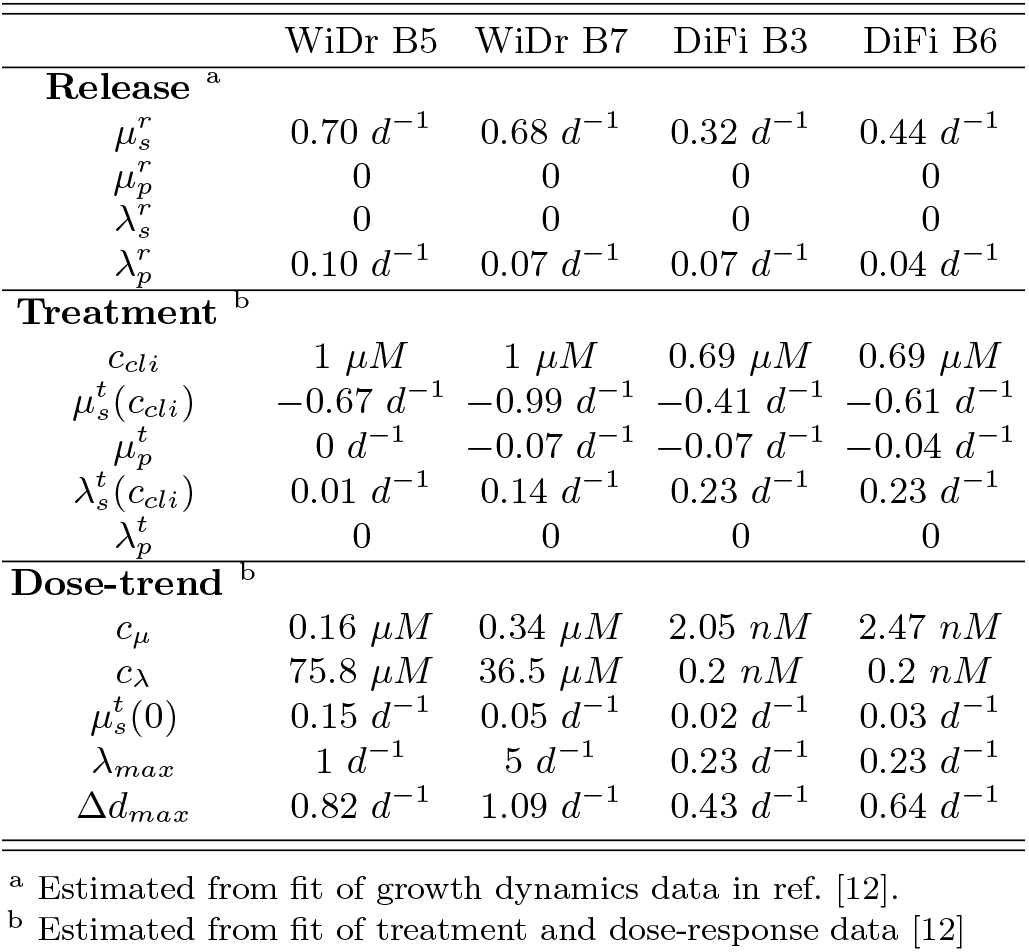
Model parameters for four cloned CRC cell line populations *WiDr cl. B5*,*B7* (treated with cetuximab and dabrafenib) and *DiFi cl. B3*,*B6* (treated with cetuximab). Parameter values are derived from experimental data from ref. [12] and shown in Supplementary Fig. 1 of this work. All parameters are expressed in units of days^−1^. Dose-dependent parameters are given at the clinically-relevant concentrations *c*_*clinical*_ = 1 *μM* dabrafenib for WiDr clones and *c*_*clinical*_ = 0.69 *μM* cetuximab for DiFi clones.

|  | WiDr B5 | WiDr B7 | DiFi B3 | DiFi B6 |
| --- | --- | --- | --- | --- |
| **Release**<sup>a</sup> |  |  |  |  |
| $\mu_s^r$ | $0.70 d^{-1}$ | $0.68 d^{-1}$ | $0.32 d^{-1}$ | $0.44 d^{-1}$ |
| $\mu_p^r$ | 0 | 0 | 0 | 0 |
| $\lambda_s^r$ | 0 | 0 | 0 | 0 |
| $\lambda_p^r$ | $0.10 d^{-1}$ | $0.07 d^{-1}$ | $0.07 d^{-1}$ | $0.04 d^{-1}$ |
| **Treatment**<sup>b</sup> |  |  |  |  |
| $c_{cli}$ | $1 \mu M$ | $1 \mu M$ | $0.69 \mu M$ | $0.69 \mu M$ |
| $\mu_s^t(c_{cli})$ | $-0.67 d^{-1}$ | $-0.99 d^{-1}$ | $-0.41 d^{-1}$ | $-0.61 d^{-1}$ |
| $\mu_p^t$ | $0 d^{-1}$ | $-0.07 d^{-1}$ | $-0.07 d^{-1}$ | $-0.04 d^{-1}$ |
| $\lambda_s^t(c_{cli})$ | $0.01 d^{-1}$ | $0.14 d^{-1}$ | $0.23 d^{-1}$ | $0.23 d^{-1}$ |
| $\lambda_p^t$ | 0 | 0 | 0 | 0 |
| **Dose-trend**<sup>b</sup> |  |  |  |  |
| $c_\mu$ | $0.16 \mu M$ | $0.34 \mu M$ | $2.05 nM$ | $2.47 nM$ |
| $c_\lambda$ | $75.8 \mu M$ | $36.5 \mu M$ | $0.2 nM$ | $0.2 nM$ |
| $\mu_s^t(0)$ | $0.15 d^{-1}$ | $0.05 d^{-1}$ | $0.02 d^{-1}$ | $0.03 d^{-1}$ |
| $\lambda_{max}$ | $1 d^{-1}$ | $5 d^{-1}$ | $0.23 d^{-1}$ | $0.23 d^{-1}$ |
| $\Delta d_{max}$ | $0.82 d^{-1}$ | $1.09 d^{-1}$ | $0.43 d^{-1}$ | $0.64 d^{-1}$ |

<sup>a</sup> Estimated from fit of growth dynamics data in ref. [12].
<sup>b</sup> Estimated from fit of treatment and dose-response data [12]

Mathematically, the cancer cell population is represented by the vector 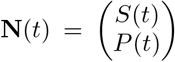 , where *S*(*t*) and *P* (*t*) *P* (*t*) correspond to the number of susceptible and persister cancer cells at time *t*, respectively. The population undergoes *q* cycles of treatment and release, with treatment duration *τ*_*t*_ and release duration *τ*_*r*_. During each phase (treatment or release), the time evolution of the population is governed by the first-order differential equation:

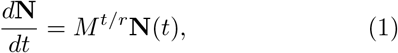

where *M* ^*t/r*^ is the transition matrix for the treatment (*t*) and release (*r*) phases, defined as,

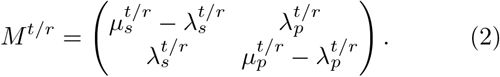

Here, 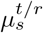 and 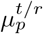 represent the net growth rates of susceptible (*S*) and persister (*P* ) cells during the treatment or release phases, while 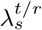 and 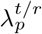 are the transition rates between the two phenotypes (Fig. 2C). Specifically, 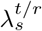 is the rate of transition from susceptible to persister cells (the “persistence transition”), and 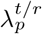 is the rate from persister to susceptible cells (the “switch-back transition”).

The predicted dynamics depend on eight key model parameters 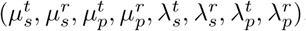. Additionally, since treatment parameters may vary with drug concentration (e.g., 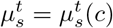), the model’s predictions are influenced by how these parameters change with dose. As a result, the model can predict a wide range of behaviors, making the dynamics quite complex.

To make the model more manageable and target it to an experimentally relevant region of the parameters, we fixed the parameters and their dose-dependencies based on the experimentally observed dynamics in our recent study of colorectal cancer (CRC) cell lines treated with clinically relevant single or combination therapies [12]. This allows us to focus on the results within this context, while also providing a framework for future studies where these parameters can be experimentally determined.

#### Drug-induced dose-dependent transition to persistence

To accurately capture the observed drug-induced and dose-dependent transition to persistence in our CRC data [12], we defined the switching rates as 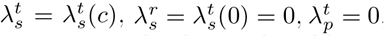, and 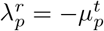 . We also note that the drug-induced hypothesis implies the initial condition *P* (0) = 0 at the beginning of the first treatment cycle. We further model the dose dependence of the switching rate using a saturating function,

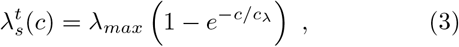

where the value of *c*_*λ*_ is derived from measurements in [12]. As illustrated in Supplementary Fig. 1 of this study, the transition-to-persistence dose dependence for DiFi and WiDr cells is accurately modeled by a monotonically increasing saturating function. In both cases, the transition rate is zero without drug, confirming that the transition is drug-induced. DiFi cells exhibit a constant transition rate for all cetuximab concentrations experimentally tested, compatible with Eq. (3) for (*c* ≫ *c*_*λ*_), while WiDr cells show a linear increase in transition rate at dabrafenib concentrations that were experimentally tested. Such linear trend is retrieved in Eq. (3) for (*c* ≪ *c*_*λ*_). The different behavior in these cell lines, despite the same functional dependence on drug concentration, reflects reflect the different treatments as well as the distinct genetic backgrounds, which influence sensitivity to treatments. For example, WiDr cells with a BRAF mutation only respond to the combination of cetuximab and dabrafenib (with clinical concentrations of 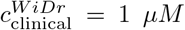 dabrafenib + 50 *μg/mL* cetuximab), while DiFi cells, which are RAS/RAF wild-type, respond to cetuximab alone (with clinical concentrations of 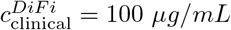 cetuximab).

##### Dose-dependent death dynamics

To model population dynamics with varying drug concentrations, we describe the dose dependency of the death rate of susceptible cells, 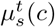. From our CRC data [12], this dependency is well captured by the equation

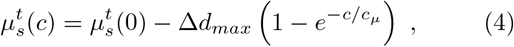

where *c*_*μ*_ is derived from measurements in [12]. The minimum inhibitory concentration (MIC), defined as the drug concentration at which the proliferation rate becomes zero, is obtained by solving 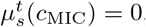, leading to

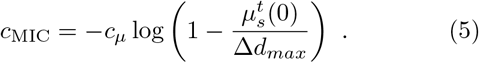

This condition will be important when integrating the model with patient pharmacokinetic data. Above the MIC, the proliferation rate becomes negative, indicating that susceptible cells die on average, with the death rate increasing with drug concentration but eventually saturating. We note that the dose-dependent behavior of both the proliferation and transition rates during treatment (illustrated in Supplementary Fig. 1), is a distinctive feature that differentiates our CRC-specific mathematical framework from previous work.

Table I reports a summary of the numerical values of the model parameters derived from data in ref. [12] and used in our analyses. For simplicity, the remainder of the main text will focus on showing the results for one representative cloned population WiDr cl. B7, leaving to the supplementary material additional information and specific insights into the behavior of the other studied cloned populations.

##### Quantification of a protocol’s long-term efficacy

Having specified the framework, our goal is twofold: (1) to characterize the conditions under which periodic treatments lead to the extinction of a treated population, and (2) to optimize treatment protocols for accelerating cancer extinction dynamics. Both objectives are addressed by quantifying two key observables derived from our model.

First, we evaluate the population’s “long-term fitness”, Λ, defined as the asymptotic per-cycle exponential rate of population growth or decay [5],

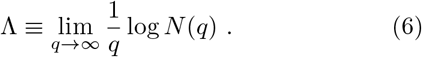

A value of Λ *<* 0 indicates population extinction (“Treatment Success”), while Λ *>* 0 corresponds to “Treatment Failure.” If Λ is slightly negative, extinction may take many cycles. The characteristic time for the population long-term dynamics is given by Λ^−1^(*τ*_*r*_ + *τ*_*t*_).

To estimate Λ from the total population number *N* (*q*), we used three indipendent methods. First, we simulate the population dynamics over many treatment-release cycles, second, we solve Eq. (1) numerically or, third, we solve it analytically,

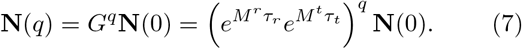

In the second case, we use the asymptotic slope of the log *N vs q* plot (Fig. 2B) as a proxy for long-term fitness, since for large *q, N* (*q*) ∼*e*^Λ*q*^, so *N* (*q*)*/N* (*q* −1) ∼*e*^Λ^. In the third case, one can show that:

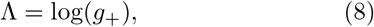

where *g*_+_ is the largest eigenvalue of the evolution matrix 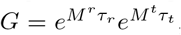 , computed via numerical diagonalization of *G* or through further analytical methods (see appendix B and refs. [3–5, 15, 20]).

In this study, we applied all three methods to evaluate long-term fitness, confirming that each approach yields the same result (Supplementary Fig. 2). Further details on the quantification of long-term fitness, including analytical calculations, are provided in Appendix B.

The second observable we used to characterize cancer long-term dynamics “asymptotic population composition”, tracks the long-term ratio of persister (*P* ) to susceptible (*S*) cells in the population,

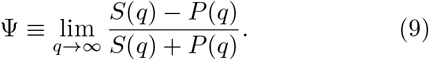

This ratio can be estimated through simulations or by solving Eq. (7). For large *q*, the asymptotic composition is given by (see appendix B)

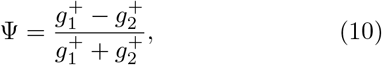

where 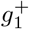 and 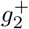 are the components of the leading eigenvector **g**^+^. A value of Ψ = −1 indicates an entirely susceptible population, Ψ = −1 indicates a persister-dominated population, and Ψ = 0 represents an equal mix of both cell types.

It is important to note that both long-term fitness and asymptotic population composition depend on the treatment protocol parameters, i.e., Λ = Λ(*τ*_*t*_, *τ*_*r*_, *c*(*t*)) and Ψ = Ψ(*τ*_*t*_, *τ*_*r*_, *c*(*t*)). This raises the question of whether a treatment protocol can be designed to optimize outcomes, such as rapidly extinguishing the cancer population. The following two sections address this question.

### B. Conditions for the design of successful intermittent protocols at constant clinical doses

To examine how treatment and release times can be effectively chosen to drive the treated cell population to extinction, we first compared periodic treatment protocols with varying release (*t*_*R*_) and treatment (*t*_*T*_ ) durations at fixed total drug exposure - keeping constant dose (*c*) per cycle and number of cycles (*q*). This approach explores whether optimizing the duration and the interval between treatment cycles can improve treatment efficacy within standard clinical frameworks, where cumulative drug dose per cycle and total treatment cycles are typically fixed [17–19].

Fig. 3 recapitulates our findings for constant-dose protocols with varying release and treatment durations at fixed clinically relevant concentrations of dabrafenib and cetuximab. We present here results for the clone WiDr B7, leaving in Supplementary Figs. 3-5 the results for the other tested CRC cell lines (WiDr B5, DiFi B3-B6), which are all consistent.

**FIG. 3.**
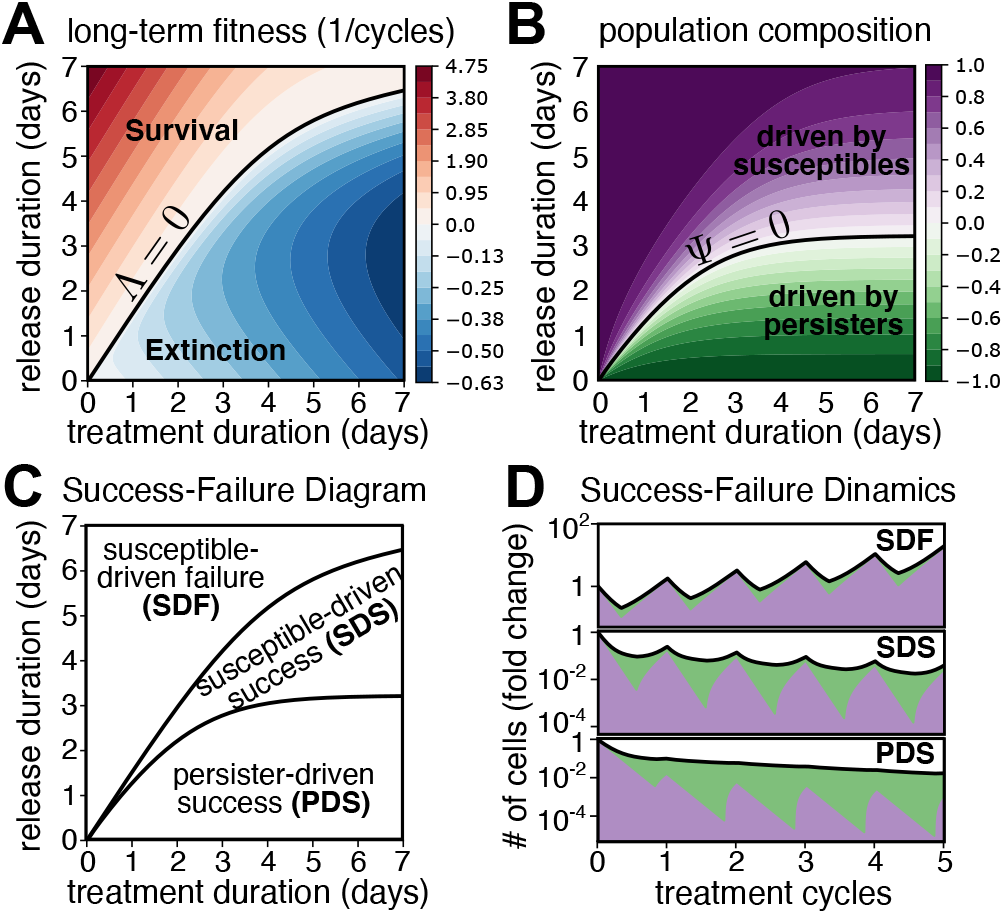
Conditions for the success and failure of intermittent treatment protocols with constant doses. **(A)** Heat-map showing long-term fitness values, Λ (unit 1/cycles), for periodic treatment protocols with varying treatment and release times on a 0 −7 day scale and drug concentration fixed to the clinical reference value *c*_*cli*_ = 1 *μM* . The long-term fitness was computed numerically using Eq. (8). The solid black line represents the Λ = 0 boundary, which separates extinction (blue, Λ *<* 0) from survival (red, Λ *>* 0) regions.**(B)** Heat-map showing asymptotic population composition, Ψ, values in correspondence of the same protocols as before. The asymptotic population composition was computed numerically using Eq. (10). Regions where Ψ *>* 0 (purple) indicate that dynamics is driven by a predominantly susceptible population, while regions where Ψ *<* 0 (green) indicate a predominantly persister population. The solid black line indicates the *ψ* = 0 boundary, where susceptible and persister fractions are equal. **(C)** The “Success-Failure (SF) phase diagram” recapitulating both long-term fitness and asymptotic population composition information. **(D)** Examples of the model’s predicted dynamics in the susceptible-driven failure (*τ*_*t*_ = 2 days, *τ*_*r*_ = 4 days), the susceptible-driven success (*τ*_*t*_ = 5 days, *τ*_*r*_ = 4 days) and the persister-driven success (*τ*_*t*_ = 5 days, *τ*_*r*_ = 1 days) regions of the protocol’s space. WiDr cl. B7 parameters (Table I).

Fig. 3A shows long-term fitness values for different constant-dose protocols, and identifies survival (Λ *>* 0) and extinction (Λ *<* 0) regions. In these protocols, the constant dose is set equal to the clinical dabrafenib dose, as reported in Table I, and treatment and release times vary over a 1-week range (Fig. 3A). By positioning a given protocol relative to the phase boundary Λ = 0 (solid black line in Fig. 3A), which separates survival and extinction regions in the phase plane (*τ*_*t*_, *τ*_*r*_), our model predicts whether a clinically-plausible protocol will effectively eradicate the population.

Moreover, it predicts the characteristic timescale of the extinction (or survival) dynamics, quantified as the number of treatment-release cycles, log 2*/* |Λ| , or the number of days, log 2*/* |Λ| (*τ*_*r*_ + *τ*_*t*_), needed to halve (or double) the treated population. For example, a protocol where clinical doses of dabrafenib are administered for *τ*_*t*_ = 5 days every week (*τ*_*r*_ = 2 days) falls in the extinction region, with long-term fitness value Λ≈− 0.5 cycles^−1^, implying a characteristic extinction timescale of about log 2*/* |Λ| ≈1.4 cycles, or 10 days. In this case, a tumor of size 10^6^ ≈2^20^ cells would require approximately 28 cycles, or 193 days, of intermittent treatment to be fully eradicated.

Additionally, increasing the treatment duration (moving horizontally in Fig. 3A) generally decreases the longterm fitness, thus reducing the average number of cycles needed for complete extinction. For instance, if the treatment duration is extended from *τ*_*t*_ = 5 to *τ*_*t*_ = 7 days while keeping the same release duration of *τ*_*r*_ = 2 days, the long-term fitness increases to Λ ≈−0.63 cycles^−1^, which implies a characteristic timescale of approximately 1 cycle. In this case, the 10^6^-cells tumor would require around 20 cycles and 180 days to be eradicated. However, it is important to note that while the characteristic timescale in cycles may decrease, the total time does not necessarily decrease, since the duration of each treatment-release cycle also increases with treatment time.

Interestingly, changing the release duration, while keeping the dose and treatment duration fixed (moving vertically in Fig. 3A), leads to more complex scenarios. This can result in either improved or worsened outcomes, reflecting a “reentrant” behavior in the extinction region (Fig. 3A). This behavior highlights a trade-off in choosing the release duration, which can be optimized to minimize the long-term fitness. We will discuss this optimization feature in the next section.

Fig. 3B shows the asymptotic population composition, Ψ, which captures the dynamic interplay between susceptible and persister cells under constant-dose intermittent treatments. Similar to the analysis of long-term fitness, we observe that the treatment-release phase plane divides into two distinct regions: one where susceptible cells predominantly drive the population dynamics (Ψ *>* 0), and another where the “keystone species” are instead persister cells (Ψ *<* 0). In other words, the dynamics can be piloted by both susceptible and persister cells depending on the protocol. For example, under a protocol where clinical doses of dabrafenib are administered for *τ*_*t*_ = 5 days every week (*τ*_*r*_ = 2 days), the dynamics is predominantly driven by persister cells. Indeed, the population composition value for this protocol is Ψ ≈−0.5 (Fig. 3B), corresponding to 75% persister cells (%*P* = 75%).

When the release duration increases while keeping treatment duration and drug concentration constant (moving vertically in Fig. 3B), the population dynamics shifts from being persister-driven at low release durations to being increasingly dominated by susceptible cells at higher release durations. This transition is due to the switch-back process during the release phase: at low release times, only a small fraction of persisters can switch back to the susceptible state. As the release duration increases, a larger fraction of persisters switch back to the susceptible state, resulting in a more susceptible-driven dynamic. Eventually, when the release duration is sufficiently long, all persisters present the treatment phase switch back during the release phase, leading to a completely susceptible-dominated population. Similarly, when the treatment duration is increased while maintaining constant drug concentration and release time (moving horizontally in Fig. 3B), the dynamics become increasingly dominated by persister cells. This shift reflects the transition to persistence during treatment, as longer treatment phases provide more time for susceptible cells to transition to the persister phenotype.

Integrating information from both long-term fitness and asymptotic population composition values, we define a comprehensive “Success-Failure phase diagram” (Fig. 3C), recapitulating all posible outcomes under constant-dose intermittent treatment protocols. We define three relevant regions (which are seen consistently across all studied cell lines, Supplementary Figs. 3-5). The “susceptible-driven failure (SDF)” region contains those treatment protocols that fail in eradicating the tumor population due to a surge in susceptible cell growth, predominantly during release intervals (top region in Fig. 3D). On the other hand, successful extinction can rely on both susceptible (“susceptible-driven success”, SDS, region in Fig. 3D) and persister cells (“persister-driven success”, PDS, region in Fig. 3D). As previously noted, short release intervals favor eradication via elimination of persister cells, whereas longer intervals shift the extinction dependence to susceptible cells. We finally note that, in principle, a fourth “persister-driven failure (PDF)” region, where survival dynamics is predominantly driven by persister cells can also exist, although not present in Fig. 3. This regime appears considering DiFi cl. B6 parameters, as reported in Supplementary Fig. 6. Interestingly, this region is narrowly located in correspondence of short treatment and release durations (0-2 days), making it a rather unstable regime, as small variations in both release and treatment time disrupt it.

### C. Combination therapies modulating persister dynamics lead to differential treatment outcomes

The previous section showed a critical role of persister cells in determining treatment outcomes. Building on this concept, we figured that combination therapies incorporating drugs specifically targeting persister cell dynamics could enhance treatment efficacy and potentially accelerate tumor eradication. To explore this idea, we conducted a proof-of-principle analysis using our quantitative framework. This analysis compared the long-term efficacy of hypothetical combination therapies aimed at targeting cancer persistence with the outcomes achieved using single-drug regimens at clinical doses of cetuximab and dabrafenib (Table I).

We considered two potential mechanisms of action for the auxiliary drug: (1) reducing the transition-to-persistence rate (type I) or (2) increasing the death rate of persister cells (type II). These mechanisms were incorporated into our model as a percentage decrease or increase in the respective rates (Supplementary Fig. 7A-B). For instance, when a molar concentration, *c*^*′*^, of a type I auxiliary drug is added, we assumed that the transition-to-persistence rate switch to a value

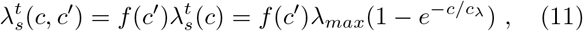

with *f* (*c*^*′*^) ∈ (0, 1). Alternative modeling choices are possible, such as modifying the characteristic dose-scale *c*_*λ*_, but both approaches ultimately decrease 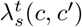. Given the absence of specific data to favor one mechanism over another, we adopted the simpler form described in Eq. (11). For subsequent analyses, we assumed an illustrative value of *f* (*c*^*′*^) = 0.5 (Supplementary Fig. 7A-B). However, we acknowledge that a more comprehensive analysis would require precise experimental characterization of *f* (*c*^*′*^). We simulated the model before and after introducing the auxiliary drug and calculated the differences in long-term fitness between single-drug and combination therapy protocols. We varied treatment and release times within a 0-7 day range as in previous analyses, keeping the drug dose constant at clinical reference values.

Interestingly, our analysis revealed distinct long-term fitness outcomes depending on whether a type I or type II drug was used (Supplementary Fig. 7C-D). Specifically, adding drugs that increase persister cell mortality (type II) generally reduced long-term fitness across protocols (Supplementary Fig. 7D), with a varying degree of efficacy. In contrast, combination therapies targeting the transition-to-persistence rate (type I) showed both beneficial and detrimental effects, depending on the specific treatment protocol (Supplementary Fig. 7C).

We integrated these findings into a phase diagram (Supplementary Fig. 7E), recapitulating the long-term fitness effects of combination therapies across protocols with varying treatment and release durations, while maintaining fixed clinical doses. Remarkably, our analysis divides the (*τ*_*t*_, *τ*_*r*_) phase plane into three distinct regions (labeled “case 1,” “case 2,” and “case 3.” in Supplementary Fig. 7E), showing different responses to combination therapies. Specifically, protocols lying in the region where susceptible cells dominate the long-term dynamics (case 1, comparing with Fig. 3) show no significant benefits from adding an auxiliary drug targeting persister cells, with type I drugs often having detrimental effects. Consequently, drugs increasing persister cell mortality had minimal impact, while those reducing the transition-to-persistence rate could inadvertently increase the number of susceptible cells. The top panel of Supplementary Fig. 7F illustrates the population dynamics in this region, (*τ*_*t*_ = 2 days, *τ*_*r*_ = 6, days, *c* = 1 *μM* ). On the other hand, protocols belonging to the region where persister cells predominantly drive the extinction of the cancer population (case 2, comparing Supplementary Fig. 7E and Fig. 3) largely benefit from type II drugs targeting persister cell mortality, while type I drugs have little to no effect (Supplementary Fig. 7C-D-E). In this region, interventions that increase persister cell death are particularly effective and reducing transition to persistence has little effect (see the middle panel of Supplementary Fig. 7F for an example of tthe predicted dynamics for the protocol, with parameters *τ*_*t*_ = 7 days, *τ*_*r*_ = 1, days, *c* = 1 *μM* ). Finally, we find empirically an intermediate region (case 3 in Supplementary Fig. 7E) region, where both combination therapies yield comparable beneficial effects on the long-term fitness (Supplementary Fig. 7C-D-E illustrated in bottom panel of Supplementary Fig. 7F, for parameters *τ*_*t*_ = 7 days, *τ*_*r*_ = 6, days, *c* = 1 *μM* , ).

In summary our analysis indicates that hypothetical combination therapies targeting the transition-to-persistence rate or persister cell mortality can enhance treatment efficacy very differently, and their impact should depend heavily on the specific treatment protocol. The next section returns to single-drug protocols to further explore optimal treatment design.

### D. The most effective treatment protocols optimize release time and drug concentration jointly

Having understood the conditions underlying successful treatment protocols, as well as the effects of adding persister-targeting drugs, we now address the problem of determining the best possible (single-drug) treatment strategy, in terms of minimizing the long-term fitness (thus minimizing residual tumor population and accelerating extinction dynamics). Mathematically, this is an optimization problem over the space of all possible intermittent treatment protocols parametrized by (*τ*_*t*_, *τ*_*r*_, *c*). The optimal treatment strategy is defined as the protocol 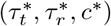 in correspondence of which the long-term fitness is minimum:

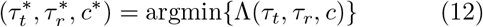

We numerically solved Eq. (12) in the range of parameters [0, 7]days *×* [0, 7]days *×* [10^−8^, 10^−4^]*M* , obtaining the best strategy

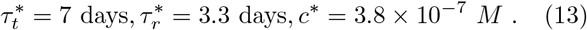

Consistently with our previous observations, we note that the best choice for treatment duration is the maximum achievable value. This is because, as we already discussed, the general effect of treatment duration is to decrease the per-cycle long-term fitness, that is minimize the average number of treatment-release cycles needed to extinguish the cancer population. On the other hand, the optimal values for the release time and drug concentration are less trivial and require further investigation. The remainder of this section focuses on discussing the optimization of these two parameters. Again, we initially consider parameters from the clone WiDr cl. B7, where a clear dependency on drug concentration of both proliferation and transition rates is seen (Supplementary Fig. 1), leaving the discussion of all other studied CRC cell lines to the end of this section.

To investigate in further detail how varying drug concentration and release duration affects the long-term efficacy of treatment protocols, we set a constant treatment duration *τ*_*t*_ = 7 days (the optimal value set by the boundary in our exploration). As described in the previous section, we computed long-term fitness and population composition varying release durations and drug concentrations to reconstruct treatment outcomes in the release time - drug concentration phase plane (Fig. 4A-B, Supplementary Fig. 11A). Our analysis reveals again that the release time-drug concentration phase plane can be divided into three regions: susceptible-driven failure (SDF), susceptible-driven success (SDS), and persister-driven success (PDS), as shown in Supplementary Fig. 11A-B. Increasing release time shifts the population towards more susceptible cells, as longer release durations allow more persister cells to switch back and proliferate. In contrast, higher drug concentrations increase the fraction of persister cells by both accelerating the elimination of susceptible cells and inducing more persisters, resulting in a persister-dominated population.

**FIG. 4.**
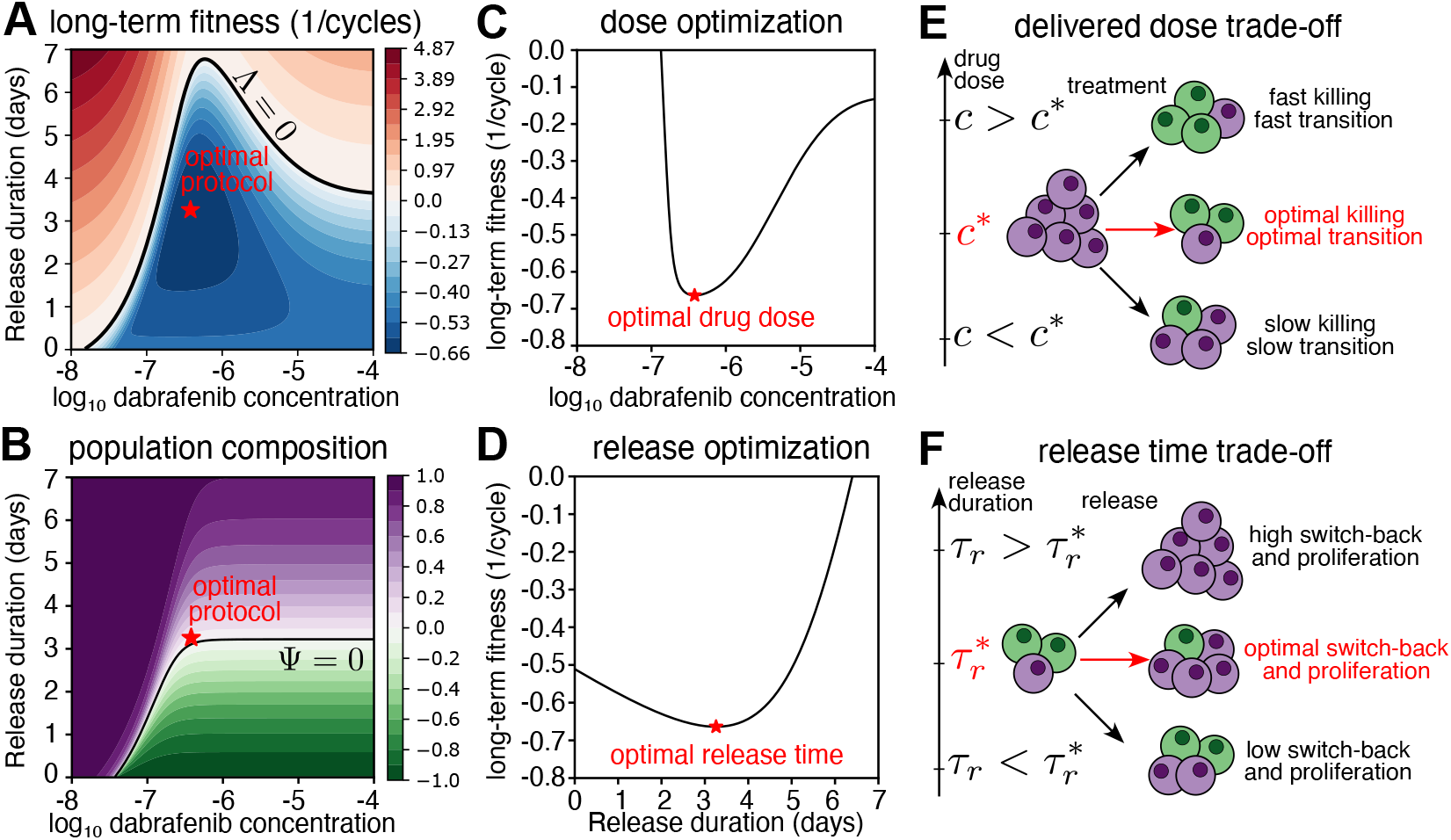
The optimal treatment protocol minimizes long-term fitness through joint optimization of release time and drug concentration. **(A-B)** Heat-maps showing long-term fitness (A) and asymptotic population composition values (B) as a function of drug concentration and release duration for a fixed treatment time of 7 days. Optimal protocol (red star) is achieved at values 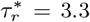 days and *c*^∗^ = 3.8 *×*10^−7^ *M* . The colorbar in panel A reports values of Λ in unit of 1*/*cycles. **(C)** The long-term fitness as a function of dabrafenib concentration is minimal (achieving a value of ≈−0.66 cycles^−1^) at the optimal drug concentration *c*^∗^ = 3.8 *×*10^−^7 *M* . **(D)** Long-term fitness as a function of release duration is minimum at the optimal time of 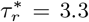 days. **(E)** Illustration of the dose-dependent trade-off between proliferation and persistence rates, defining an optimal drug concentration. **(F)** Illustration of the interplay between resumed proliferation and subsequent drug efficacy, indicating an optimal release time: if the release time exceeds this value, a larger fraction of persister cells reverts to a vulnerable state, enhancing the effect of the next cycle. Conversely, if too short, most treated cells remain in a persister state and are unaffected by subsequent treatment. Both too short and too long release times yield higher final population sizes than the optimal release time. WiDr cl. B7 parameters (table I).

Fig. 4A shows that increasing release duration as well as drug concentration affects long-term fitness. For protocols with fixed treatment and release times while varying drug concentration (moving along horizontal lines in Fig. 4A), long-term fitness initially decreases at low drug concentrations, reaching negative values around the clinical dabrafenib concentration (*c* = 1 *μM* ), before increasing again. Similarly, for protocols with fixed treatment duration and drug concentration while varying release time (moving along vertical lines in Fig. 4A), long-term fitness first decreases, then increases, eventually reaching positive values for sufficiently high release durations. This behavior suggests the presence of an optimal protocol (red star in Fig. 4A), which minimizes long-term fitness by jointly optimizing drug dose and release time. Fig. 4B shows how these findings are paralleled by population composition. For the specific WiDr B7 CRC cloned population used here, the optimal protocol balances the fractions of persister and susceptible cells (red star in Fig. 4A-B). This balance in persister and susceptible cell dynamics is crucial for achieving optimal extinction outcomes. However, this feature is not universal (and thus is likely to be patient-specific), as different parameters may lead to varying optimal population compositions, as seen in Supplementary Figs. 8-10, which show results for other CRC clones.

Fig. 4C shows the dose-dependence of long-term fitness at the optimal release time 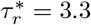 days, where the optimal drug concentration is *c*^∗^ = 3.8 *×*10^−7^ *M* , less than half the clinical reference value (*c* = 1 *μM* ). This finding suggests that lower drug doses may reduce toxicity and improve therapeutic efficacy by increasing extinction speed.. This dose optimization is a consequence of the dabrafenib dose-dependent behavior of the WiDr cell line, as both death and transition-to-persistence rates increase monotonically with drug concentration (depicted in Supplementary Fig. 1 and recapitulated by Eq. (3) and Eq. (4)). There is a trade-off between using lower drug concentrations, where both death and transition rates are low, and higher doses, where both rates are fast (Fig. 4E). As shown in our previous study [12], the optimal concentration balances this trade-off by maximizing killing dynamics while maintaining a moderate transition rate to persistence. Suboptimal concentrations occur when lower doses fail to increase the death rate sufficiently, while higher doses overly increase the transition rate with little additional effect on death rate, which is already at its maximum (Fig. 4E).

Similarly, Fig. 4D shows the long-term fitness dependence on release time at the optimal drug concentration *c*^∗^ = 3.8 *×*10^−7^ *M* . We note that, within clinically relevant concentration ranges, which lie around the optimal concentration value, neither significantly shortening nor excessively extending the release time yields optimal outcomes, suggesting a precise balance between dosing intervals. This timing-based optimization highlights the interplay between persister cell reversion to a vulnerable state and subsequent drug efficacy. As shown in Fig. 4F, if the release time is extended, more persister cells revert to a drug-sensitive state, enhancing the population’s response in the following cycle. However, if the release time is too short, most treated cells remain in a persister state, reducing the overall impact of the next cycle.

We replicated the analyses discussed so far with the other CRC cloned populations. Supplementary Fig. 8-10, show that release time optimization is found in both our CRC cell lines across both cloned populations, underscoring the importance of precisely timed drug administration to maximize therapeutic efficacy. On the other hand, dose-dependency of the persister transition is specific of WiDr clones treated with different doses of dabrafenib, hence resulting in a optimal dose minimizing the long-term fitness (for both WiDr B5 and B7). Conversely DiFi populations (both B3 and B6) do not show an increased transition rate to persistence for increasing drug concentrations, leading to a plateau of optimal concentration values for long-term fitness (Supplementary Fig. 9-10C)

Finally, we note that the optimization features discussed in this section are neither intuitive nor immediately apparent, highlighting the power of mathematical models in guiding the design of optimal treatment protocols. To illustrate this, we compared the efficacy of the predicted optimal protocol, defined by (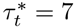 days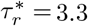 days, *c*^∗^ = 3.8 ×10^−7^ M), with that of an alternative protocol characterized by: (i) a higher drug dose, *c*^*′*^ = 10^−5^ M *> c*^∗^, and (ii) continuous drug administration, where the waiting time between consecutive treatments is zero (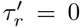 days 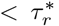). Counterintuitively, our model predicts that administering a lower drug dose while allowing longer waiting periods between treatments results in higher efficacy. To reinforce this prediction, we simulated several stochastic trajectories (Appendix D of the treated cell population dynamics under both the optimal and suboptimal protocols (Supplementary Fig. 11C) and we computed the corresponding extinction time distributions (Supplementary Fig. 11D). The results confirm that the optimal protocol, despite involving less frequent and lower-intensity treatments, outperformed the suboptimal one, reducing the average extinction time by more than 20% (73 days for the optimal protocol versus 93 days for the suboptimal protocol), confirming the value of quantitative modeling in uncovering counterintuitive strategies. The next section uses our quantitative framework to explore the efficacy of continuous and intermittent dosing strategies in scenarios that take into account measured pharmacokinetics parameters.

### E. Optimal intermittent dosing of dabrafenib rivals the efficacy of continuous treatments

To further assess the potential clinical applicability of our quantitative framework, we seeked to connect it to the ongoing clinical debate surrounding continuous versus intermittent dosing regimens [19]. To this end, we transitioned from the simplified assumption of constant drug treatment profiles to explore models reflecting the dose kinetics observed in patients treated with similar doses of the same targeted drugs we used with our CRC cell lines (Table II). This required shifting from a constant concentration to a time-dependent profile that decreases after each administration. We assumed that the reduction in drug concentration directly correlates with decreased death rates and transition rates for susceptible cells, as measured for our cell lines.

**TABLE II.** Adopted CRC cell lines and treatments [12].

| CRC cell lines | Sensitivity | Treatment |
| --- | --- | --- |
| DiFi <sup>a</sup> B3-B6 | EGFR blockade | cetuximab ($100 \mu\text{g/mL}$) |
| WiDr <sup>b</sup> B5-B7 | EGFR blockade<br>BRAF inhibition | cetuximab ($50 \mu\text{g/mL}$)<br>dabrafenib ($1 \mu\text{M}$) |

<sup>a</sup> RAS/RAF wild-type
<sup>b</sup> BRAF V600E-mutated

Inspired by experimental observations [18], we model the in-patient dose profile as a double exponentially decreasing function of the form:

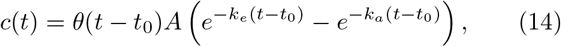

which, albeit very simple, recapitulates known key aspects of drug pharmacokinetics, such as temporal delays (*t*_0_), maximum induced plasma concentration (*c*_*max*_), and both dose absorption (rate *k*_*a*_) and elimination (rate *k*_*e*_) dynamics (Fig. 5A).

**FIG. 5.**
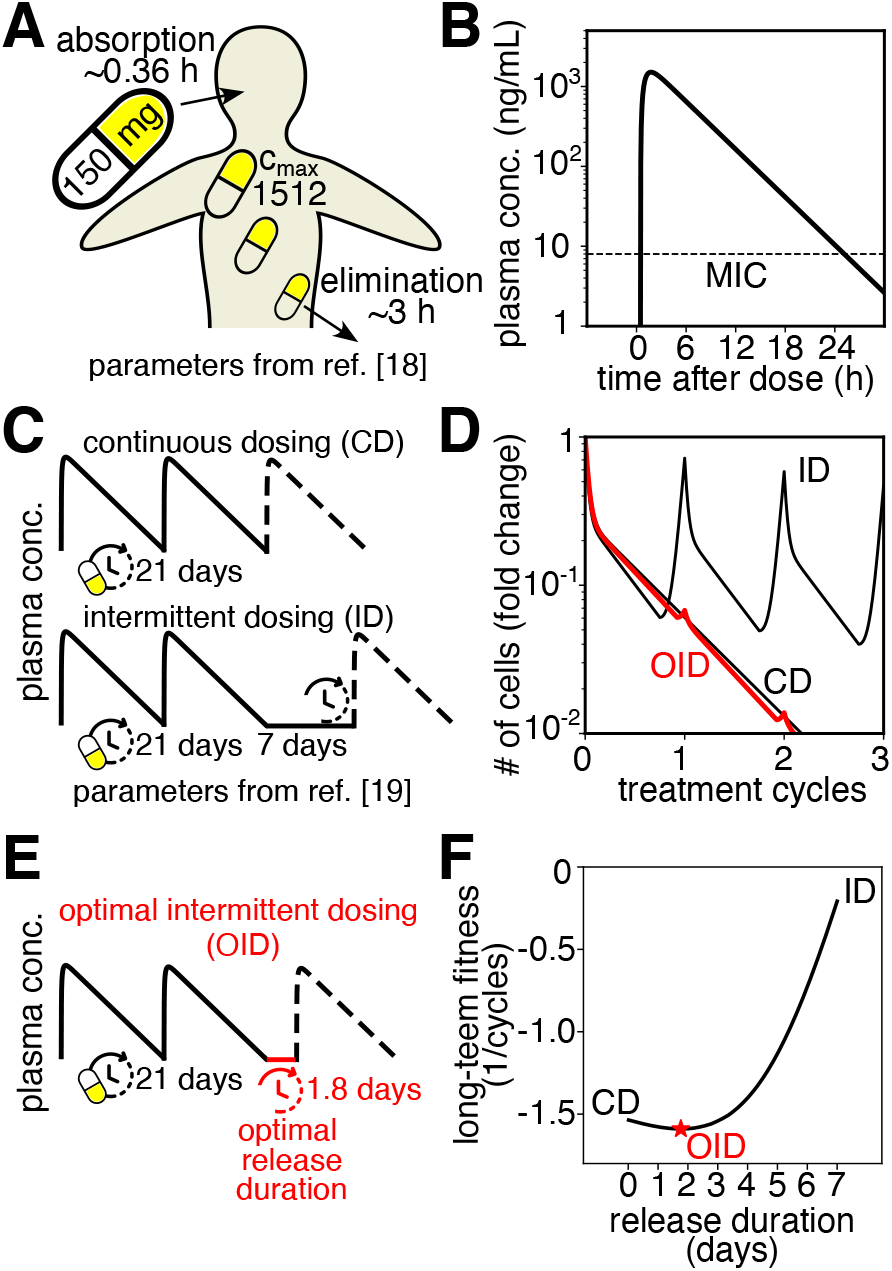
Exploiting drug pharmacokinetics to optimize intermittent dosing of dabrafenib in patients. **(A)** Illustration of the typical time course of drug concentration in patients, comprising absorption and elimination dynamics. Reported values for dabrafenib dose (150 *mg BID*), peak plasma concentration (*c*_*max*_ = 1512), absorption (*k*_*a*_ = 0.36*h*^−1^) and elimination (*k*_*e*_ = 3 *h*) rates, have been derived from study [18] and used to fix the plasma concentration profile reported in panel B. **(B)** Inferred plasma concentration profile, showing a rapid absorption phase ( ≈ 0.36 *h*), followed by a slower exponential decline of drug concentration over time ( ≈ 3 *h*). After 1 day the plasma concentration reaches the minimum inhibitory concentration of dabrafenib ( ≈ 8 *ng/mL*, dashed horizontal line). **(C)** Constant versus intermittent dosing of dabrafenib, as recently compared in clinical trial [19]. **(D)** Comparison of predicted cancer decline for constant and intermittent dosing of dabrafenib. Red line shows dynamics under the optimal intermittent dosing strategy. **(E)** Illustration showing optimal intermittent dosing parameters. **(F)** Long-term fitness as a function of release time showing non-optimality of both continuous and intermittent dosing strategies considered in study [19]. Optimal release duration, defining the optimal intermittent strategy, is equal to *τ*_*r*_ = 1.8 days (red star)

Importantly, we used the experimental measurements from the clinical trial in ref. [18], characterizing the population pharmacokinetics of dabrafenib, to fix the parameters in Eq. (14). Specifically, we use reported values for the absorption rate, *k*_*a*_ = 1.88 h^−1^ (*τ*_*a*_ = log(2)*/k*_*a*_ ≈ 0.36 h), maximum induced (peak absorption) plasma concentration, *c*_*max*_ = 1512 ng mL^−1^, lag time, *t*_0_ = 0.48 h, and area under the curve, *AUC* = 8820 ng mL^−1^ h to estimate *A* and *k*_*e*_ as (see appendix C for further details)

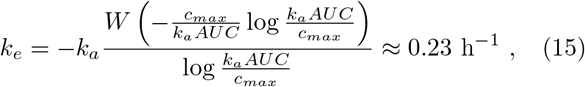

with *τ*_*e*_ ≈ 3 h, and where *W* is the Lambert function, and

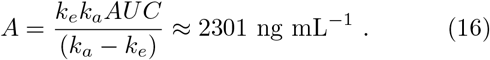

The resulting doseprofile predicted for a patient,

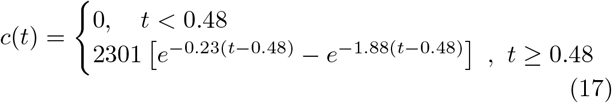

is plotted in Fig. (5B) over a daily time span. We note that in about one day, the inferred dabrafenib plasma concentration reaches the minimum inhibitory concentration (MIC), 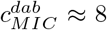 ng mL^−1^ (dashed horizontal line in Fig. 5B), as estimated from Eq. (5). Below the MIC, dabrafenib concentrations are no longer capable of inducing susceptible cell death, leading us to set 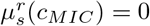 in our model, and motivating the need of renovating dabrafenib administration on a daily time scale [18, 19]. Furthermore, we note that the time dependence of the drug concentration makes the dose-dependent parameters in our model (susceptible cells’ death and transition-to-persistence rates), dependent on time. Supplementary Fig. 12 shows the specific time dependence of our model parameters under these conditions.

Equipped with these additional features, we simulated our model to investigate the problem of whether intermittent dosing of dabrafenib might be preferable to constant dosing regimens in a patient. We note that a recent clinical study [19] addressing this same question reported no evidence in favor of an intermittent treatment strategy. Here, by comparing the model’s predicted cancer dynamics under the same continuous and intermittent dosing protocols considered in [19], depicted in Fig. 5C, we revised this comparison from the more quantitative point of view of assessing the long-term dynamics of the treated cell population. For simplicity, we computed the long-term dynamics under both continuous and intermittent dosing protocols further approximating Eq. (17) by considering an instantaneous absorption dynamics.

Consistently with the findings of the clinical-trial [19], an intermittent dosing (ID) strategy where dabrafenib is administered for 21 days every 28 days (release time of one week) shows inferior performance in our model compared to a continuous dosing (CD) strategy in which dabrafenib is administered every day (Fig. 5D). Interestingly, however, when we exploit our model’s flexibility to compute how long-term fitness changes as a function of the release time interval between consecutive dabrafenib administrations, we see that neither the continuous dosing (CD), nor the intermittent dosing (ID) considered in the clinical trial in ref. [19] are predicted to be optimal choices, when aiming at minimizing the long-term fitness of the treated cell population. Instead, we find that an intermittent protocol with a release time interval of about 2 days, 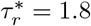 days (Fig. 5E), minimizes the long-term fitness (Fig. 5F), yielding the fastest achievable extinction dynamics of the total cancer population (red line in Fig. 5D). Thus, our analysis leads us to surmise that an optimized intermittent protocol, particularly when its release timing is finely tuned, may be able to yield outcomes that not only rival but may even exceed those of a continuous dosing regimen.

## III. DISCUSSION

Targeted therapies based on EGFR inhibitors like cetuximab, and their combination with dabrafenib for BRAF-mutated tumors, were shown to extend median survival by a few months in colorectal cancer patients when added to standard chemotherapy, compared to chemotherapy alone [21]. This is a modest but non-negligible outcome, and it is important to understand how it can be improved upon. While a limitation to the efficacy is certainly due to emergence of resistance, phenotypic transitions and persistence also likely play a role [8, 11, 22]. Given the limitations of current targeted therapies and the potential role of mechanisms beyond resistance, we explored the efficacy of periodic treatment-release protocols. In this analysis, we have assumed that resistance does not develop. While of course we acknowledge that resistance is a significant clinical challenge, we note that recent work suggests that even in the presence of resistant clones, treatment release periods can still be beneficial if competitive non-resistant clones remain [20]. Additionally, in the case of resistance emergence due to drug-induced mutagenesis, mathematical modeling has also shown that less aggressive treatments could potentially lead to improved outcomes [23]. However, while these lines of evidence point to the idea that periods of treatment release can generally be an advantage, they also show that such release period need to be precisely targeted on the specific parameters of a tumor population (and adapted if these parameters change), or they risk to be detrimental. Since the precise knowledge of these parameters in a clinical setting is scarce, the practical applicability of these concepts remains very limited, although as personalized-medicine assays advance, one could envisage that such parameters may be used to optimize treatment time-administration protocols, together with choice of treatments (see below).

To avoid these complications, we have rooted our analysis on observations and parameter values measured in CRC cell lines [12]. Importantly, we note that the parameters and behaviors were different in the two tested CRC cell lines and treatments, supporting the idea that patient specificity might be relevant. Under the radical (but necessary) assumption that the parameters measured on cell lines may guide us to reveal relevant scenarios, our findings suggest a non-intuitive landscape for the effectiveness of periodic treatments in controlling and potentially eradicating treated cancer cell populations with persisters. Specifically, we show how long-term fitness defines the boundaries between successful therapy and treatment failure and how population composition in susceptible *vs* persisters is essential for understanding optimal treatment protocols, due to a delicate balance between treatment (which kills susceptible cells but produces persisters) and release phases (which clear persisters but allow susceptible cells to proliferate). A consequence of this complex balance is the observation of reentrant behavior in extinction dynamics (Fig. 3), where increasing the duration of release periods can counterintuitively optimize the eradication process, by providing sufficient time to clear persisters (a result unattanable under continuous treatment). In scenarios where treatment fails, unchecked proliferation of susceptible cells during release intervals prevails. Conversely, effective treatment regimens capitalize on either efficient elimination of susceptible cells or control of persister generation, achieving optimal outcomes when they can accomplish both. This insight underscores a delicate interplay dependent on parameters such as the persister transition rates that may be challenging to access in realistic situations, where slight adjustments could be critical in tipping the balance between therapy failure and success. Our framework also allows for exploring hypothetical combination therapies targeting persister death and the transition from susceptible cells, which can significantly improve the outcome of periodic treatment protocols (Supplementary Fig. 7).

If drug dose impacts on both the death rate of susceptible cells and their transition rate to persisters, as in WiDr cells treated with dabrafenib and cetuximab, the scenario is even more complex, due to the quantitative trade-off between effective killing and persister production. As shown previously [12], an increasing dose ramp decreases persister production in a single treatment compared to the same total dose administered continuously. While this work focuses on constant alternated treatments, we speculate that combining optimal dose ramps with optimized protocols could further improve patient outcomes if persisters are relevant. For periodic treatments at constant doses, we identified an optimal dose near the clinically relevant dose of dabrafenib, where the trade-off between proliferation suppression and persistence induction yields the best outcomes. Collectively, these findings suggest that specifically targeting the persistence mechanism could enhance therapeutic efficacy.

We recognize that there are significant challenges in translating these highly theoretical optimizations into practical clinical applications. Key factors such as pharmacokinetics and pharmacodynamics and effective behavior of a treated tumor *in vivo*, where several factors related to the micronvironment can play an adverse or favorable role, may play a crucial role in determining the effective dose experienced by a tumor, introducing variability that must be carefully considered [17–19]. Moreover, the fact that cetuximab is typically administered in combination with chemotherapy regimens like FOLFIRI or FOLFOX, rather than as a monotherapy, adds another layer of complexity [24–26]. Encouragingly, a recent study showed that intermittent administration of FOLFIRI and panitumumab achieves better outcomes than continuous treatment [27]. However, substantial inter-patient variability in plasma cetuximab concentrations [17] further complicates the prediction of treatment efficacy. Spatial effects, including heterogeneous drug distribution within the body, toxicity concerns, and the influence of the tumor microenvironment, must also be accounted for when considering the clinical implementation of our findings. To take a first step towards mitigating this gap of knowledge, we have simulated our model with time-dependent doses and constraints derived from pharmacokinetics data on dabrafenib [18, 19], therefore with realistic values for the time-dependent plasma concentrations of the treatment, but still assuming that the death-rate and transition-rates were those observed in cell lines. Under these assumptions, the model prediction is that treatment-release periods of about two days would still outperform continuous treatment (but much longer release times would not).

However, given the complexity of the problem and the extremely limited available knowledge, our work serves merely as a proof of concept, offering a foundation for future research to validate and refine these hypotheses. One promising approach could involve experimental comparisons of treatment regimens in cell lines, patientderived organoids, and mouse xenografts [28, 29], providing increasingly sophisticated and controlled models for assessing the efficacy of various treatment strategies.

These experimental systems would allow for testing the predictive power of our simplified modeling framework in more clinically relevant environments. Alternatively, a more empirical approach might involve directly designing clinical trials to test various time-dependent protocols in patients, such as maintaining an equal total dose while varying the administration schedule [17–19]. Such trials could help verify the broader hypothesis that “keystone” persisters may be relevant for therapy outcomes, and could be guided by mathematical models based on in vitro systems and measured pharmacokinetic parameters. If successful, these protocols could ideally extend beyond colorectal cancer (CRC) to other cancers, such as lung cancer treated with osimertinib, where persister transitions have been observed in vitro [7, 30]. However, to motivate these protocols for other cancer treatments, a more advanced quantitative in vitro characterization of persister behavior, similar to that performed for CRC, would be essential [12]. This approach, combining theoretical insights with empirical validation, may effectively reach conclusions within a more reasonable timeframe.

In conclusion, these insights offer a roadmap for the rational design of personalized treatment schedules based on precise measurements and mathematical modeling, and suggest that a deeper understanding of the dynamic interplay between drug action, phenotypic plasticity, and recovery phases during treatment release could help isolate optimal conditions for treatment success, to inform more effective therapy strategies.

## ACKNOWLEDGMENTS

This work was supported by Associazione Italiana per la Ricerca sul Cancro, AIRC IG Grant no. 23258. MC was supported by an AIRC fellowship for Italy - id. 28177. SP was supported by Fondazione Umberto Veronesi. We thank Edo Kussell, Mariangela Russo, Alberto Bardelli, Ciro Mercurio, Andrea Bertotti and Silvia Marsoni for feedback and useful discussions.

## AUTHOR CONTRIBUTIONS

M.C.L. and S.P. conceived the study and contributed with key ideas at different stages. M.C. and A.D.B. performed data analysis, model simulations and analytical calculations. M.C., M.C.L. and S.P. wrote the paper. All authors read and approved the final version of the paper.

## DATA AND CODE AVAILABILITY

All data and code used in this study will be made openly available on GITHUB.

## Appendix A: Considered colorectal cancer cell lines

In [12], the population dynamics of two microsatellite-stable (MSS) human colorectal cancer (CRC) cell lines were analyzed: the RAS/RAF wild-type “DiFi” and the BRAF V600E-mutated “WiDr”. These cell lines were treated with specific drug regimens: cetuximab alone at a dose of 100 *μ*gmL^−1^ for DiFi, consistent with its approval for metastatic CRC patients lacking RAS and BRAF mutations, and a combination of cetuximab (50 *μ*gmL^−1^) with the BRAF inhibitor dabrafenib (1 *μ*M) for WiDr, reflecting its potential effectiveness in BRAF-mutant CRC cases. For both models, individual clones were isolated, and their growth and drug sensitivity closely matched those of the parental populations.

In this study, we evaluated our model in four different parameter configurations, derived from fitting population dynamics data for four cloned populations: WiDr cl. B5, WiDr cl. B7, DiFi cl. B3, and DiFi cl. B6, based on analyses in [11, 12].

To determine the parameters of untreated sensitive cells, we measured growth rates using growth assays under standard cell-culture conditions. For parameters under drug treatment, instead, we analyzed data from two types of drug-response growth assays. The first was a dose–response assay, where CRC clones were exposed to increasing concentrations of targeted therapies. This assay was used to evaluate growth curves, defined as the number of live cells over time and drug concentration, and to quantify the interplay between cell growth and the transition to a persister state during drug treatment. The second was a single-dose assay, involving three weeks of exposure to a constant drug concentration. This assay revealed a biphasic killing curve, characterized by an initial rapid decline in the number of sensitive cells, followed by a slower phase indicative of persister emergence, resembling patterns observed in bacterial populations. The surviving persister fraction showed a gradual but measurable decline over time, suggesting a slow tendency for persisters to die.

All treatment data and the model fits are presented in Supplementary Fig. 1. Inferred model’s parameters are summarised in Table I.

## Appendix B: Estimation of long-term fitness and asymptotic population composition

This section discusses different equivalent ways of evaluating the treated population’s long-term fitness:

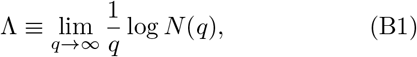

and asymptotic composition:

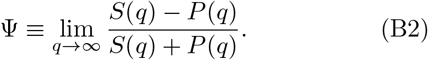

### Method 1: Estimation via dynamics simulations

One first possibility for estimating the long-term fitness, exploiting the actual definition of this parameter, is to simulate the dynamical system of Eq. (1) over many treatment-release cycles and then compute the long-term fitness as the asymptotic slope of the *logN* vs *q* plot:

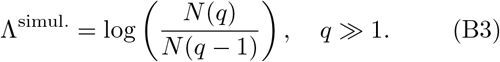

Indeed, in the long-term limit *q* ≫1, *N* (*q*) ∝ *e*^Λ*q*^, *N* (*q* −1) ∝ *e*^Λ*q*^*e*^−Λ^, so that *N* (*q*)*/N* (*q* −1) ∝ *e*^Λ^. Asymptotic population composition is straightforwardly obtained from the definition, Eq. (B2), as

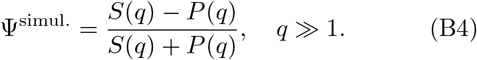

We used the scipy.integrate library to integrate the dynamical system of Eq. (1) over many treatment-release cycles. We then used Eq. (B3) and Eq. (B4) to evaluate long-term fitness and asymptotic population composition values.

### Method 2: Estimation via numerical diagonalization

This method considers the solution of the dynamical system of Eq. (1), that is Eq. (7), and aims to find the eigenvalues *g*_+_, *g*_−_ and eigenvectors 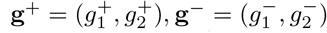 of the evolution matrix matrix 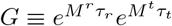 to write **N**(*q*) = *G*^*q*^**N**(0) in the form

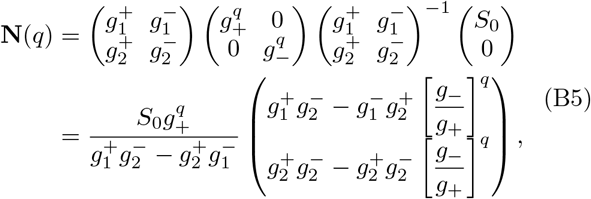

where we used the drug-induced hypothesis *P*_0_ = 0. In the long-term limit *q* ≫ 1 one has:

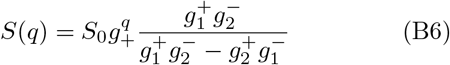

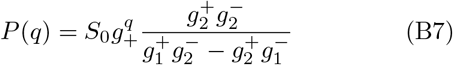

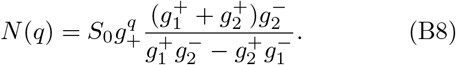

Therefore, using Eq. (B1) and Eq. (B2), one finds

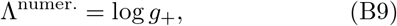

which corresponds to Eq. (8) in the main text and

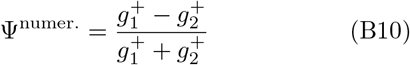

which is Eq. (10) in the main text.

We used the numpy.linalg library to compute the eigenvectors and eigenvalues of the matrix 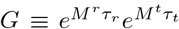, and then used Eq. (B9) and (B10) to evaluate long-term fitness and asymptotic population composition values.

### Method 3: Analytical estimation

While, numerically the diagonalization of the evolution matrix, *G*, is straightforward, analytically this constitutes a daunting task, mostly because *G* is defined via exponentiation of treatment and release matrices. Nonetheless, we will show that one can use the eigenvalues, 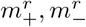 and 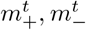 , and the eigenvectors, 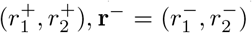 and 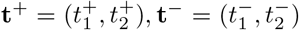, of the release and treatment evolution matrices, which can be easily computed, to reconstruct the eigenvalues and eigenvectors of *G*.

This is done by exploiting the decomposition:

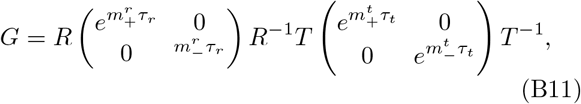

where *R*≡ (**r**^+^**r**^−^) and *T*≡ (**t**^+^**t**^−^), which, upon computation of all terms in the right-hand-side of Eq. (B11), leads to an expression of the form:

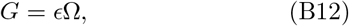

where *ϵ* is a leading exponential factor that can be factored out and Ω a residual correction matrix. This is a simplification of the problem, from the analytical point of view, because computing the eigenvalues *ω*_+_, *ω*_−_ and eigenvectors ***ω***^+^, ***ω***^−^ of Ω is now analytically feasible, yielding:

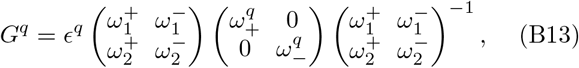

which, compared to Eq. (B5), identifies the eigenvalues *g*_*±*_ and eigenvectors, **g**^*±*^ of *G*, as:

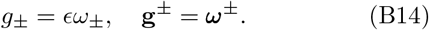

Therefore, using Eq. (B9) and Eq. (B10), we can write

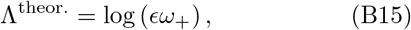

and

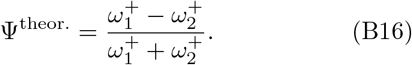

In the remainder of this section, we derive the specific expressions of *ϵ*, ***ω***^*±*^ and *ω*_*±*_ for our CRC-specific case.

First of all, eigenvalues, 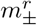 , and eigenvectors, **r**^*±*^, of the release matrix, *M* ^*r*^, satisfying the equation:

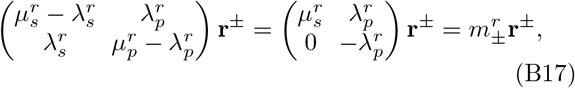

where we used the fact that in our setting 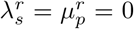, can be easily derived and read:

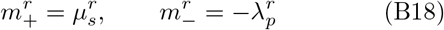

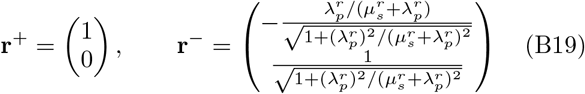

On the other hand, eigenvalues, 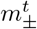, and eigenvectors, **t**^*±*^, of the treatment matrix, *M* ^*t*^, satisfying:

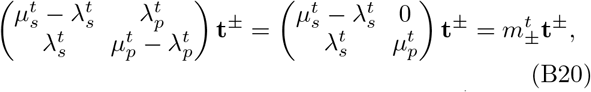

where we used the fact that in our setting 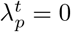, read:

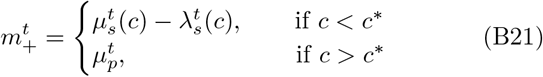

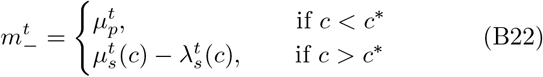

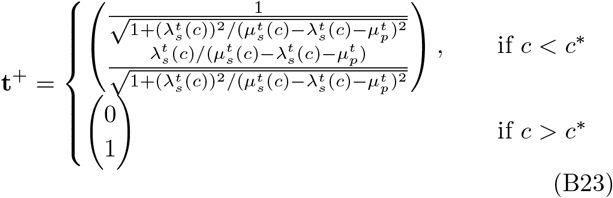

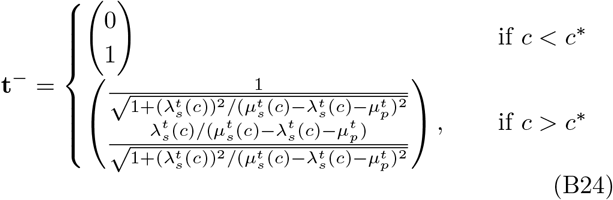

Note the switch between eigenvalues and eigenvectors when crossing the critical concentration *c*^∗^, defined as the dose concentration value at which 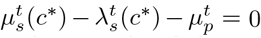 . Since *c*^∗^ is typically lower than the considered drug concentration values (for example, for the clone WiDr B7 *c*^∗^ is close to 3.8 *×*10^−8^ *M* ) we will proceed with considering the *c > c*^∗^ case.

We now compute step by step all terms appearing in Eq. (B11). For the sake of convenience, we set:

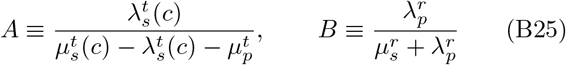

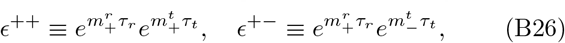

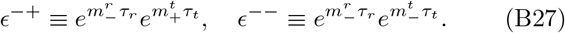

With these definitions, 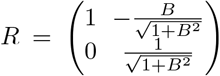 , and 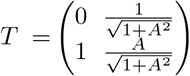 , so that we have

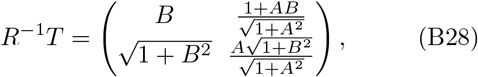

which multiplied to the right by 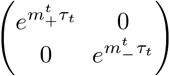, and to the left by 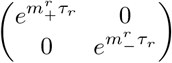 , gives

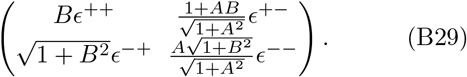

Finally, multiplication to the left by *R* and to the right by *T*^−1^, yields

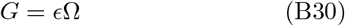

with

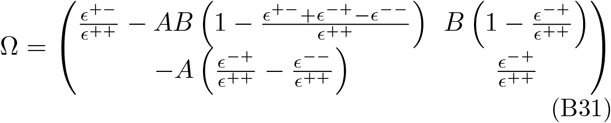

and

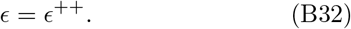

Eigenvalues and eigenvectors of Ω can therefore be computed as:

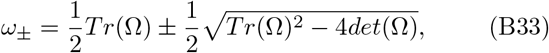

and

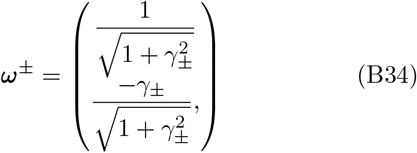

where

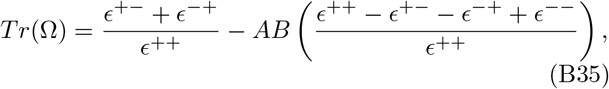

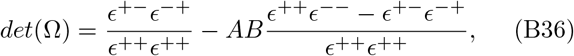

and

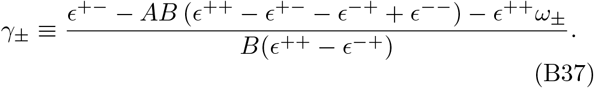

Therefore, substituting these quantities into Eq. (B15) and Eq. (B16) we find

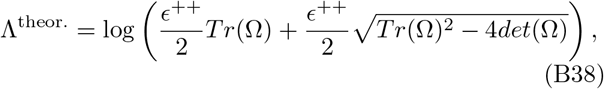

and

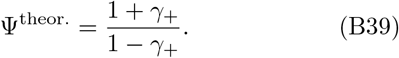

We point out that the analytical estimates derived in this section (Eq. (B38) and Eq. (B39)) are exact. Indeed, we verified that the different long-term fitness and asymptotic population composition estimation methods discussed in this section, summarised by Eqs. (B3)-(B9)-(B38) and Eqs. (B4)-(B10)-(B39), are all consistent and provide the same quantities (Supplementary Fig. 2).

### Relation with previous work

An exact analytical expression for the long-term fitness was previously derived by Patra and Klumpp in [3], using a different analytical approach. Since Eq. (B38) presented here is also exact, it can be considered an equivalent solution specific to our drug-induced scenario.

Approximated analytical solutions have also been reported in earlier studies. For instance, Kussell and coworkers in [5] provided an approximated analytical solution for long-term fitness, valid for long release and treatment durations (referred to as the “Large populations - the deterministic limit” in [5]). In our framework, an equivalent approximated solution, valid for the druginduced case where 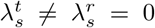 and 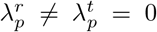, can be derived by retaining only the leading terms in either Eq. (B31) or Eqs. (B33) and (B34). Specifically, for long treatment and release durations, *τ*_*t*_, *τ*_*r*_ ≫ 1, where *ϵ*^++^ ≫ *ϵ*^+−^, *ϵ*^−+^, *ϵ*^−−^, we can approximate

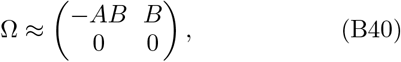

yielding *ω*_+_ ≈ − *AB*. The approximated formula for the long-term fitness therefore reads

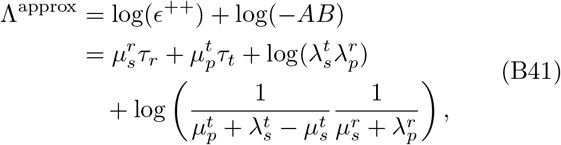

which is equivalent to Eq. (8) in [5] within our model regime. We verified the validity of this approximation by comparing it with exact estimation methods.

## Appendix C: Derivation of dabrafenib absorption and elimination dynamics observed in patients

For the absorption and elimination dynamics of dabrafenib in patients we assume a double exponential trend of the form

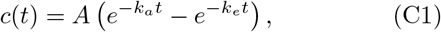

where *k*_*a*_ and *k*_*e*_ are the absorption and elimination constant rates, respectively, and *A* is a factor whose determination will be discussed below. Allowing for a lag time *t*_0_ before actual absorption of dabrafenib, the previous equation reads

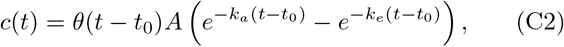

with *θ*(*t*) the standard Heaviside function. In the previous equation, *A* and *k*_*e*_ are unknowns that we now express in terms of measured experimental quantities ( *c*_*max*_, the peak absorbed dose, *AUC*(0, *τ* ), the area-under-the-curve in the experimental time window and the measured absorbed rate *k*_*a*_).

First, *c*_*max*_ can be determined as *c*(*t*^*^) where *t*^*^ is such that *c*^*′*^(*t*^*^) = 0. After simple math, one finds 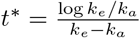, and 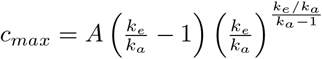, so that

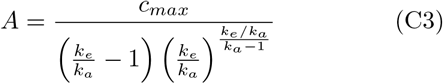

Second, from the definition of 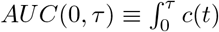 we find 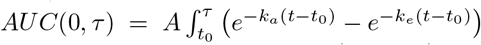 which we approximate to 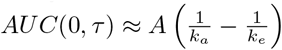, so that

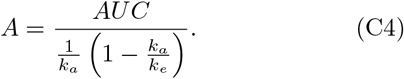

Using equations C3 and C4 we can therefore write

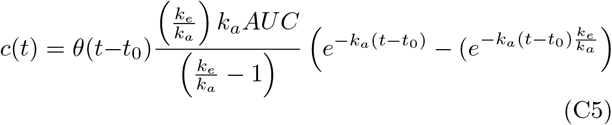

where *k*_*e*_*/k*_*a*_, solving the equation 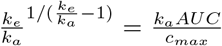 can be written as

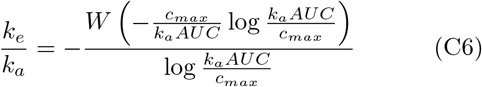

with W the lambert (product logarithm) function.

Substituting the experimental values derived from the clinical trial [18], *c*_*max*_ = 1512 *ng mL*^−1^, *AUC* = 8820 *ng mL*^−1^*h, k*_*a*_ = 1.88 *h*^−1^ and *t*_0_ = 0.48 *h*, we can finally write

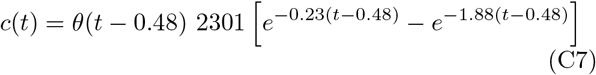

which is equation 17 in the main text and the function plotted in Fig. 5.

## Appendix D: Stochastic simulations

In this study, we built on our previous work [12] and used a coarse-grained accelerated version of the Gillespie algorithm [31, 32]. This approach groups stochastic events into discrete time intervals of fixed duration Δ*t*, reducing computational demands. The approximation assumes that events within the interval Δ*t* are independent and can be sampled from a multinomial distribution. The parameter Δ*t* was carefully adjusted to balance numerical accuracy and computational feasibility.

Stochastic trajectories of susceptible and persister cells, {*S*(*t*), *P* (*t*)}, were derived by mapping the population dynamical rates into probability transition rates that define the one-step process {*S*(*t*), *P* (*t*)}. This method uses the deterministic dynamics predicted by the dynamical system of Eq. (1) as a basis. For instance, the dynamical rate 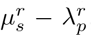, describing the time variation of susceptible cells during the release phase, is interpreted as the transition probability rate per unit time for creating a new susceptible cell, i.e., the transition *S* → *S* + 1.

Using the Gillespie algorithm, the system evolves reaction-by-reaction by sampling waiting times, *τ* , from an exponential distribution with a rate equal to the sum of all possible transition rates. After determining which reaction occurs, the system’s state is updated as *t* → *t*+*τ* and **N**(*t*) → **N**(*t* + *τ* ). While this method is exact, it can become computationally expensive when simulating large populations, as in our case. To address this, coarsegrained algorithms, which aggregate reactions over discrete time intervals, provide an efficient alternative. Several such variants exist to accelerate the Gillespie algorithm, and we employed one tailored to our system.

## Supplementary Figures for

**Supplementary Fig. 1.**
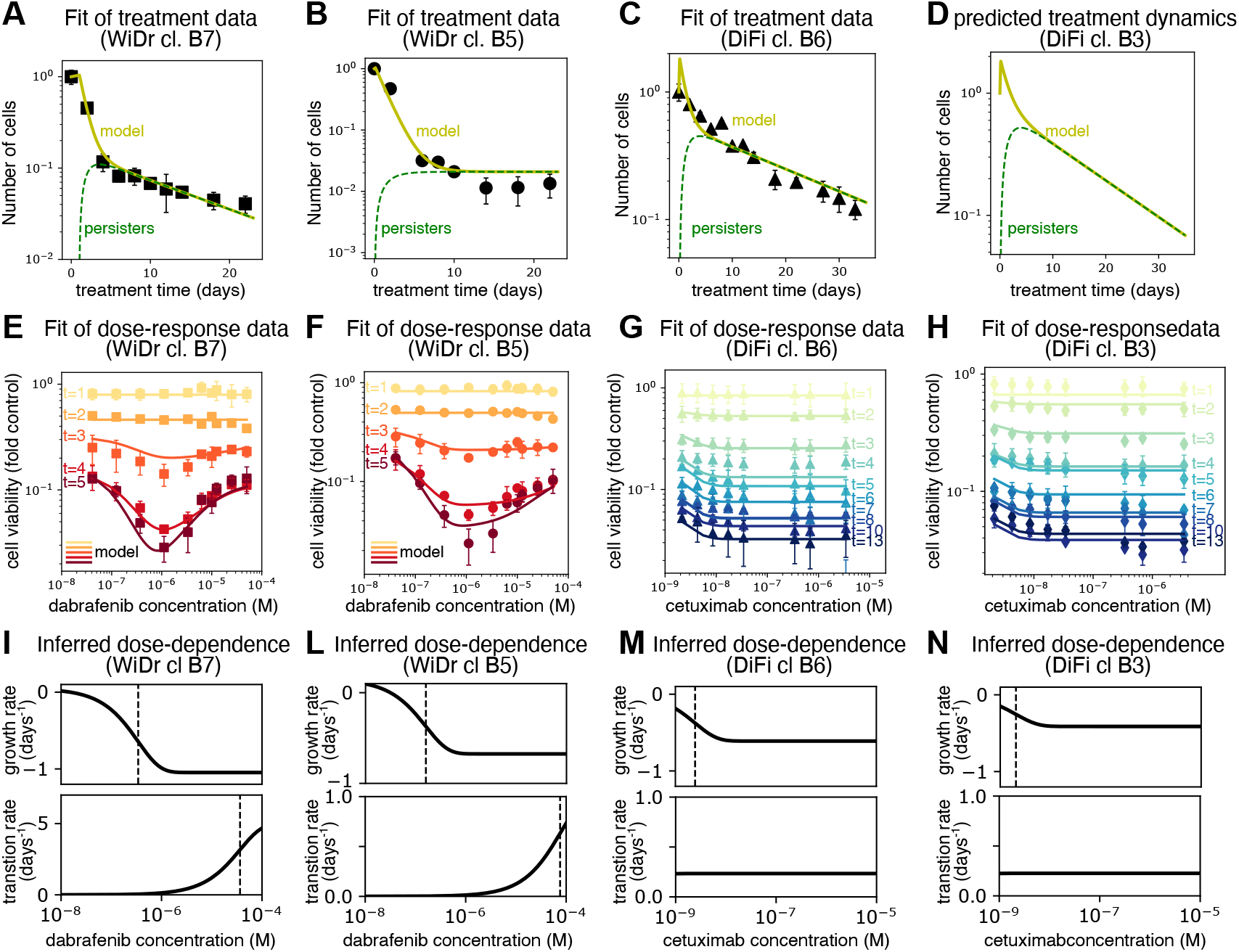
Quantitative comparison of model and experimental population dynamics of colorectal cancer cell-line data from [Russo et al. 2022] allows us to fix parameters and dose-dependencies. (A-D) Fold-change of viable cells versus time of drug exposure, assessed as discussed in [Russo *et al*. 2022]. Black symbols represent averages of biologically independent experiments. The yellow solid line indicates the fit of the model to the data, whereas the expected fraction of persister cells is represented by the dashed green line. (E-H) Experimentally measured dose-responses growth curves (points) and model fit (solid lines). The doses-response datasets were normalized to the growth of the untreated cells, as previously discussed in [Russo *et al*. 2022]. Data points are reported as mean values *±* standard deviation. The color-code represents assessment of dose-response at different timepoints. (I-N) Inferred dose-dependence of susceptible cells’ proliferation (top) and transition-to-persistence rates (bottom). Vertical dashed lines are in correspondence of the typical concentration scales, *c*_*μ*_, *c*_*λ*_ , as described in Eq. (3) and Eq. (4) of the main text. We note that within the range of drug concentrations accessible experimentally, the transition to persistence dependence is either linear (WiDr clones, panel I-L) or constant (DiFi clones, panel M, N).

**Supplementary Fig. 2.**
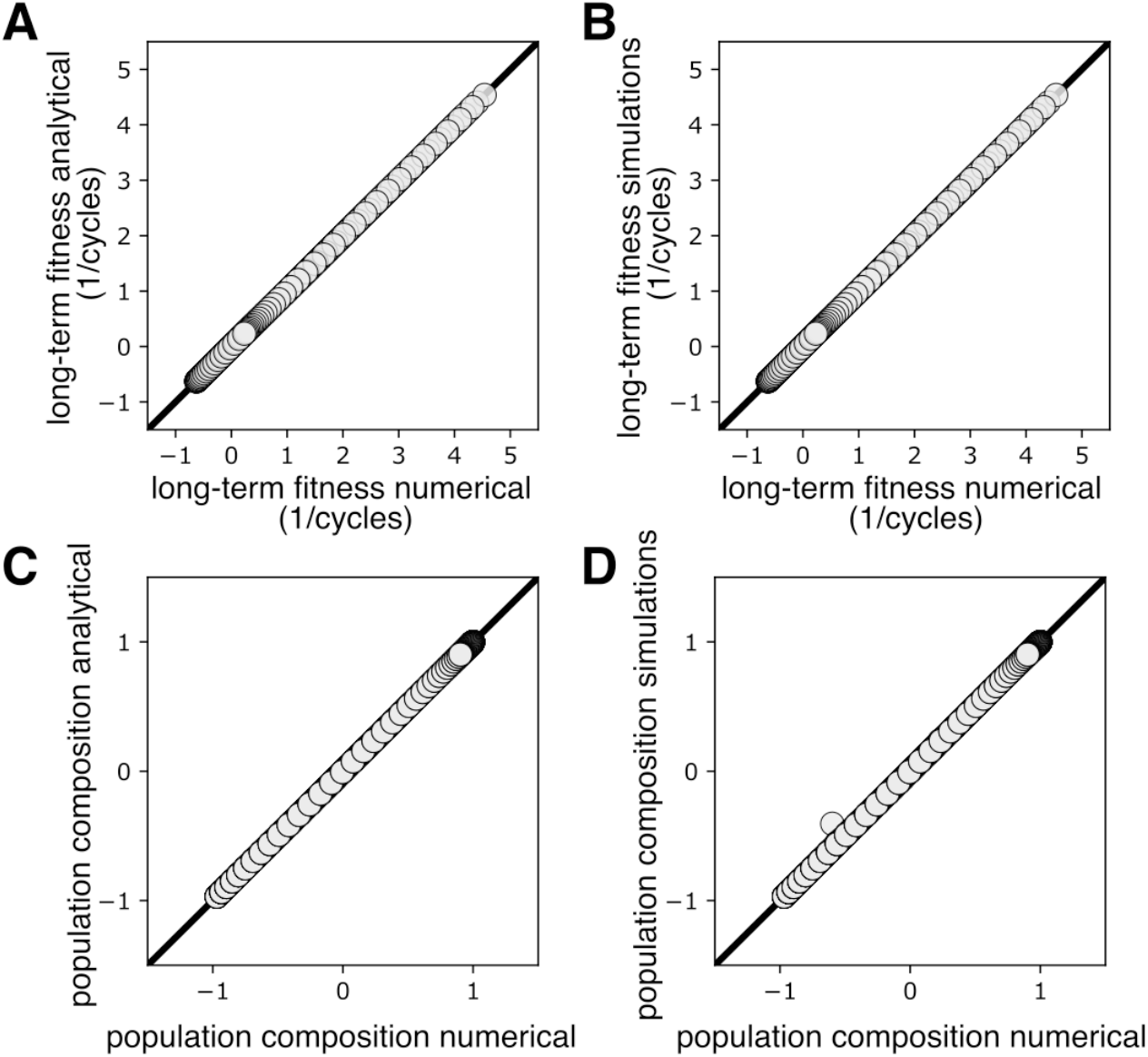
Validation of different numerical and analytical methods to obtain the long-term fitness and asymptotic population composition. (A-C) Agreement between analytical and numerical long-term fitness (A) and asymptotic population composition (C) estimation methods. (B-D) Agreement between simulations and numerical long-term fitness (B) and asymptotic population composition (D) estimation methods. Each dot corresponds to long-term fitness (population composition) estimates on a different protocol, obtained varying treatment and release durations in a 0–7-day scale, while fixing drug concentration to the clinical reference value of 1 *μM* . WiDr cl. B7 parameters.

**Supplementary Fig. 3.**
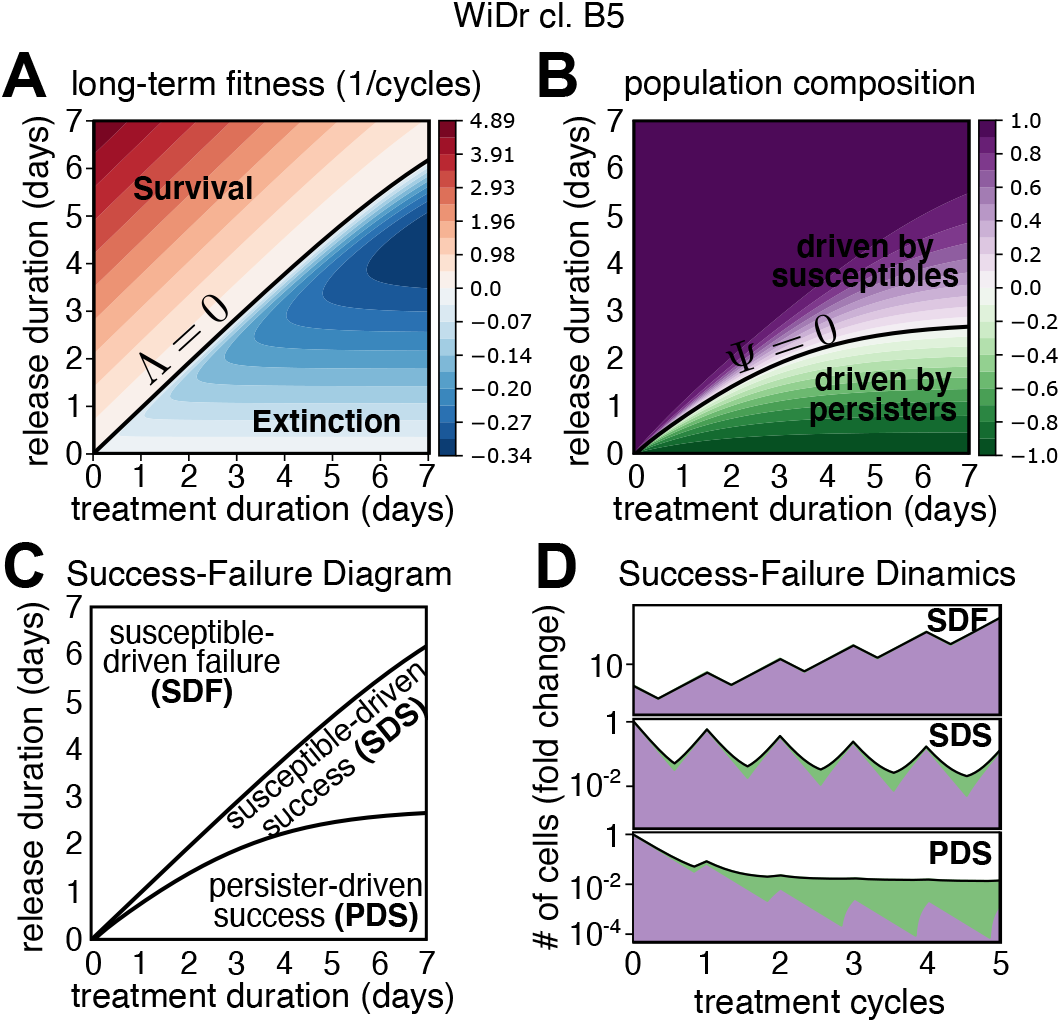
Replication with WiDr cl. B5 parameters of the same analysis reported in Fig. 3 of the main text. See caption of Fig 3. in the main text.

**Supplementary Fig. 4.**
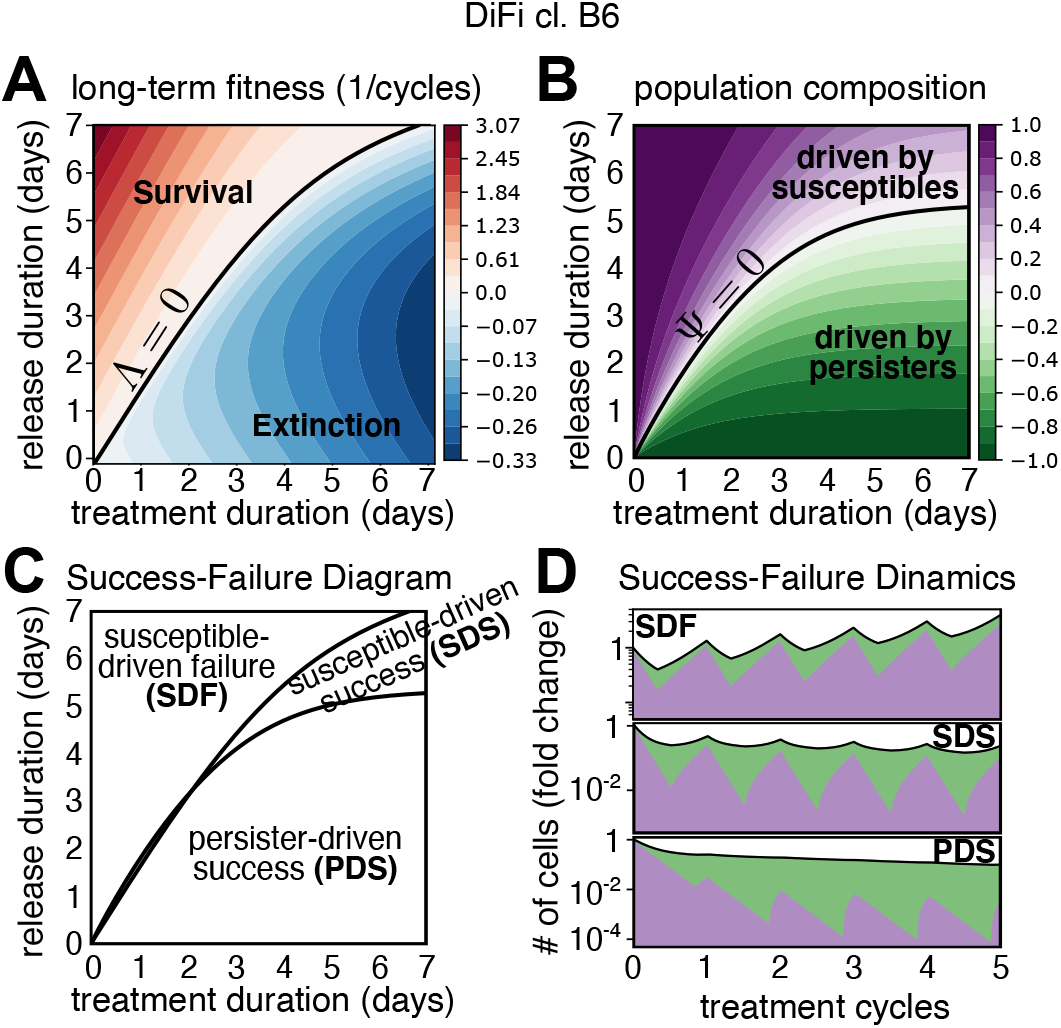
Replication with DiFi cl. B6 parameters of the same analysis reported in Fig. 3 of the main text. See caption of Fig 3. in the main text.

**Supplementary Fig. 5.**
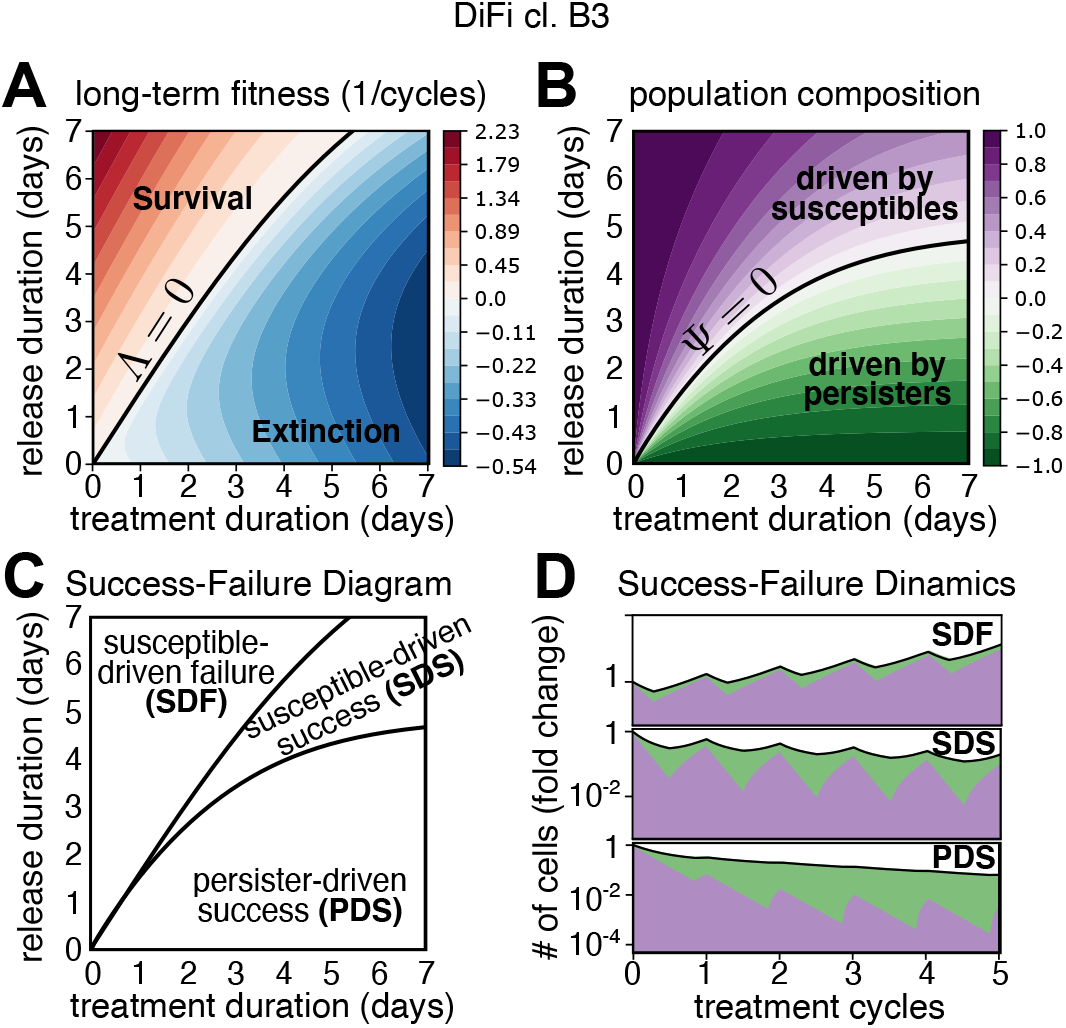
Replication with DiFi cl. B3 parameters of the same analysis reported in Fig. 3 of the main text. See caption of Fig 3. in the main text.

**Supplementary Fig. 6.**
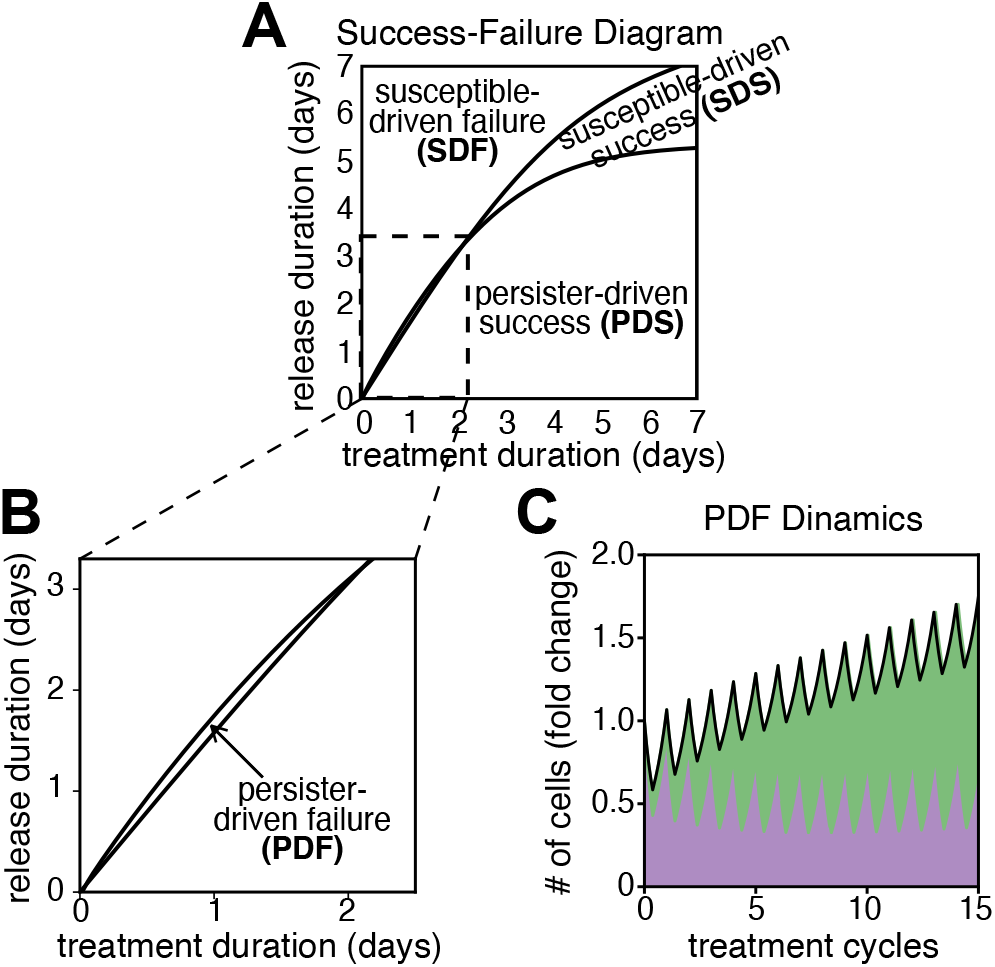
(A-B) A fourth “persister-driven failure (PDF)” region is seen in the DiFi cl. B6 clone over a short 0–3-day temporal window. (C) Model’s predicted dynamics in the “persister-driven failure (PDF)” region. Green and purple areas, represent number of persister and susceptible cells, respectively. DiFi cl. B6 parameters.

**Supplementary Fig. 7.**
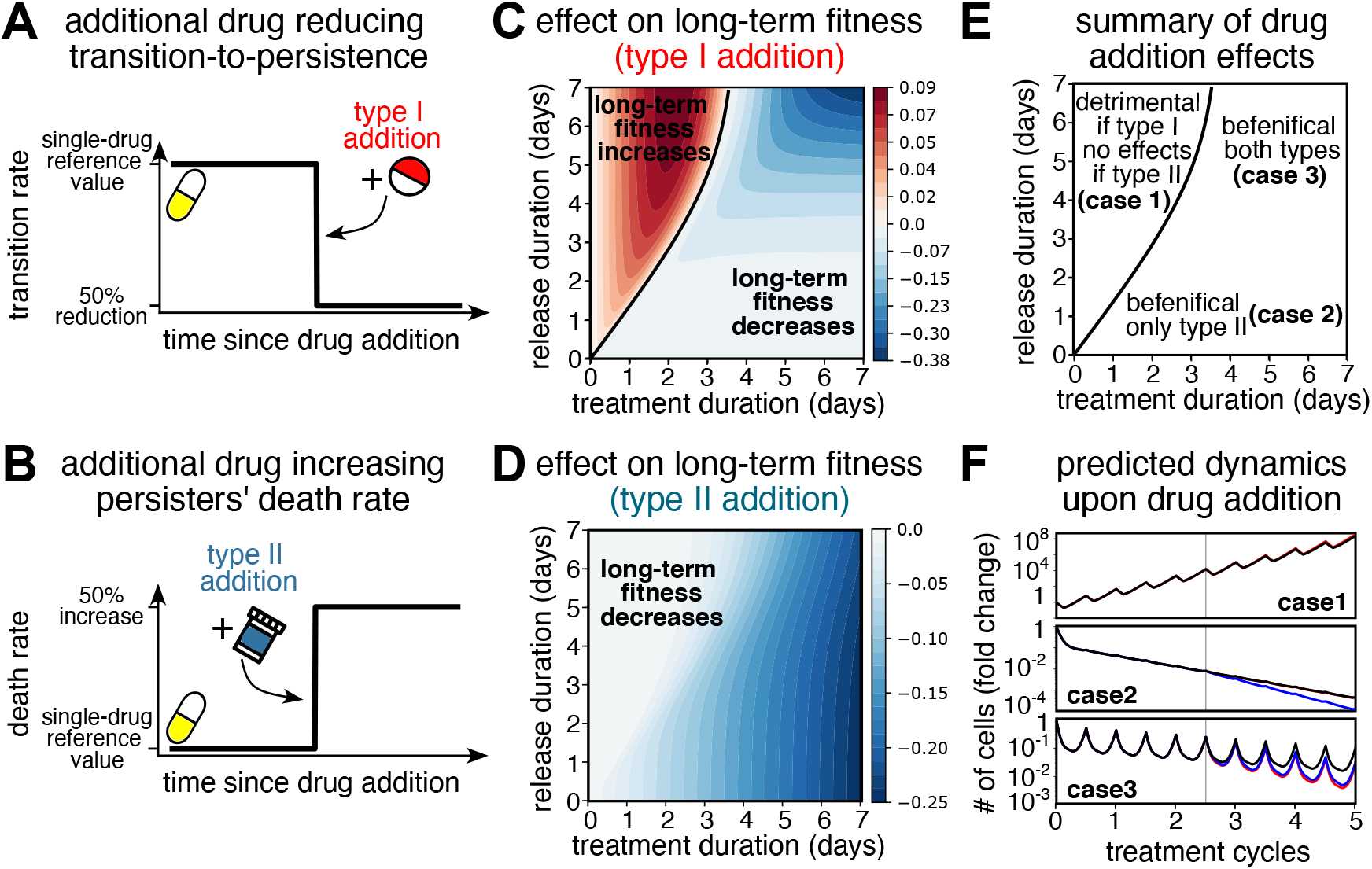
Quantification of the long-term fitness effects of combination therapies targeting persister cells dynamics. (A-B) Explored combination therapies include additional drugs that either reduce the transition-to-persistence rate (type I, panel A), or increase persisters’ death rate (type II, panel B). (C-D) Heat-maps showing the difference in long-term fitness between combination therapies, either type I (panel C) or type II (panel D), and single-drug treatment protocols with varying treatment and release times on a 0–7-day scale and drug concentration fixed to clinical reference levels. Red (blue) regions correspond to an increase (decrease) in long-term fitness upon addition of the auxiliary drug. (E) Phase diagram integrating information from panel C and D, individuating three main possible effects upon drug addition: detrimental or negligible effects (case 1), beneficial only if type II drugs are added (case 2) and beneficial under either drug (case 3). (F) Model’s predicted dynamics under protocols from each of the three cases, confirming the expected different behavior. WiDr cl. B7 parameters (see Table I in main text).

**Supplementary Fig. 8.**
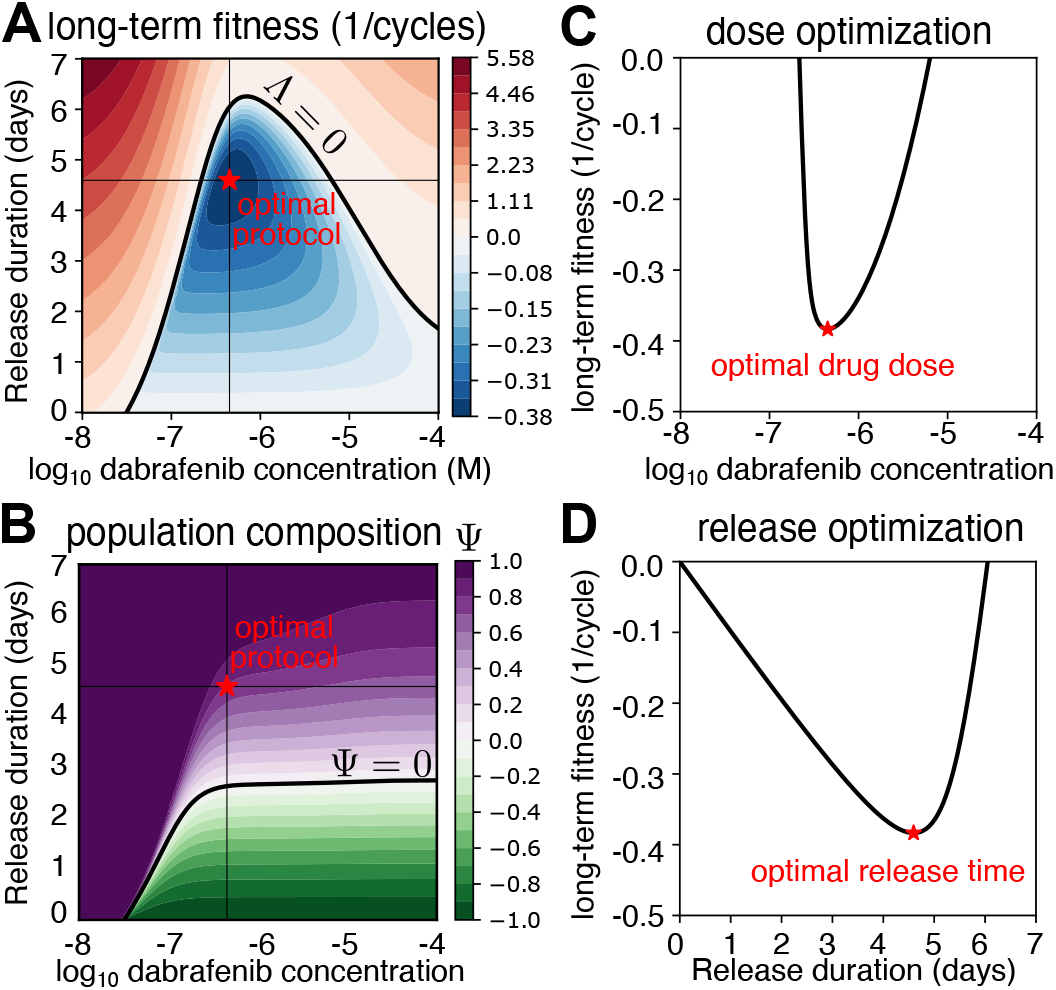
Same analysis reported in Fig. 4 of the main text with WiDr cl. B5 parameters. Optimal protocol (red star) is achieved at values (*c* = 4.5 *×* 10^−7^ *M* ,*τ*_*r*_ = 4.6 days). See caption of Fig. 4 in the main text.

**Supplementary Fig. 9.**
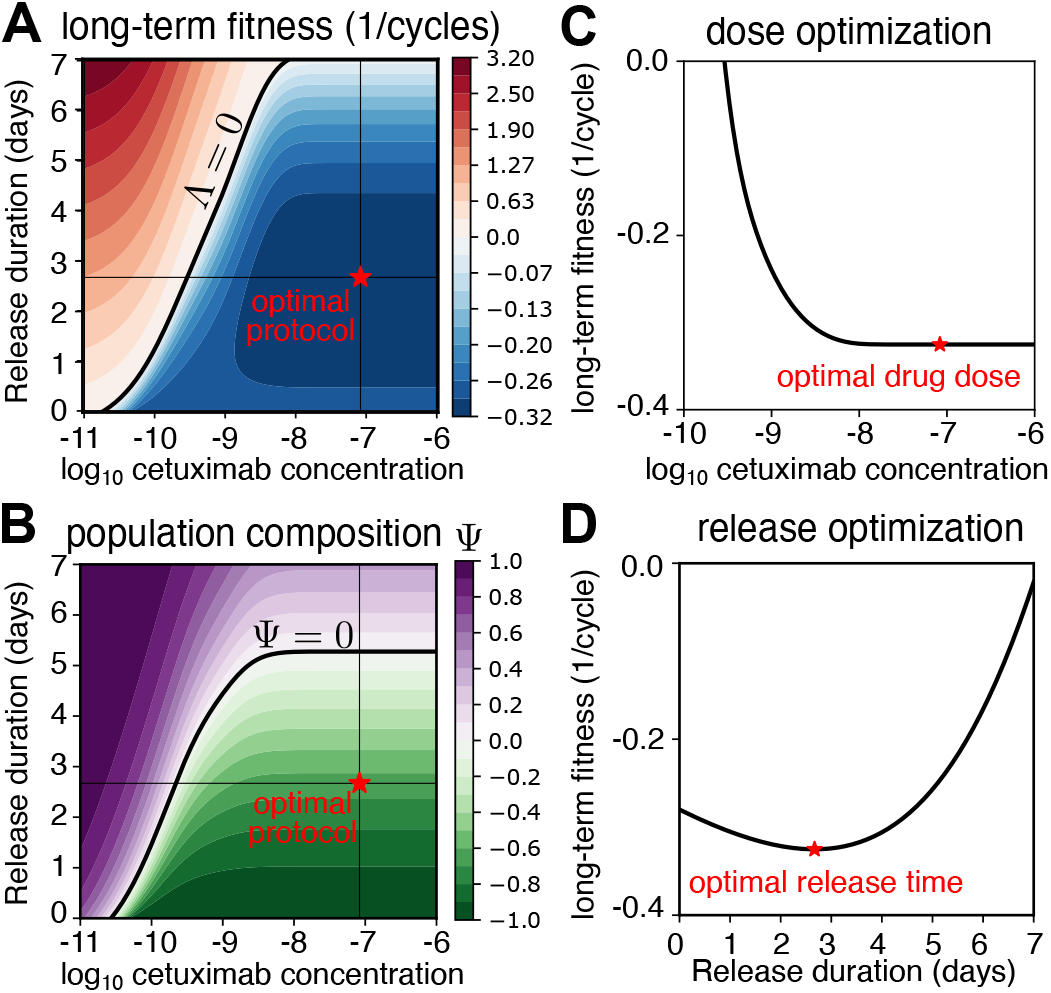
Same analysis reported in Fig. 4 of the main text with DiFi cl. B6 parameters. Optimal protocol (red star) is achieved at values (*c* = 8.3 *×* 10^−8^ *M* ,*τ*_*r*_ = 2.7 days). See caption of Fig. 4 in the main text.

**Supplementary Fig. 10.**
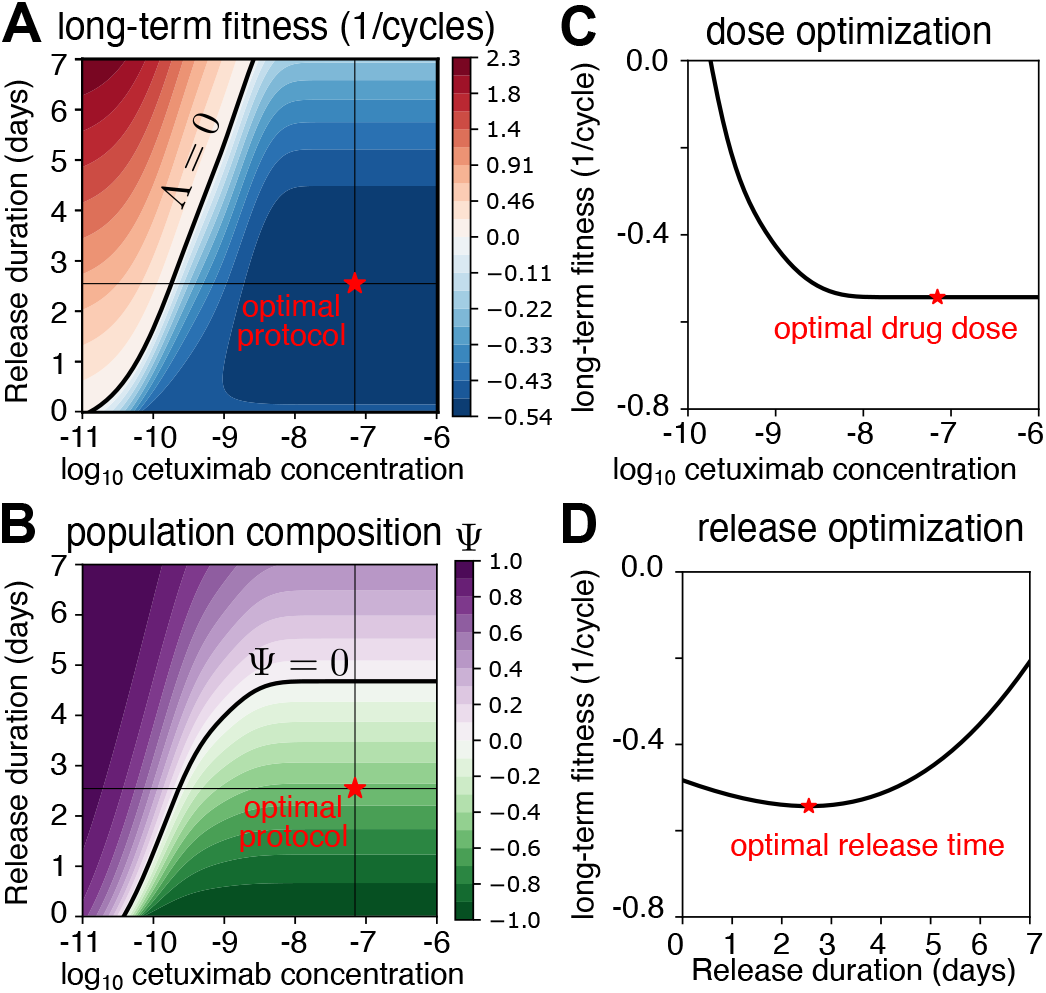
Same analysis reported in Fig. 4 of the main text with DiFi cl. B3 parameters. Optimal protocol (red star) is achieved at values (*c* = 7 *×* 10^−8^ *M* ,*τ*_*r*_ = 2.5 days). See caption of Fig. 4 in the main text.

**Supplementary Fig. 11.**
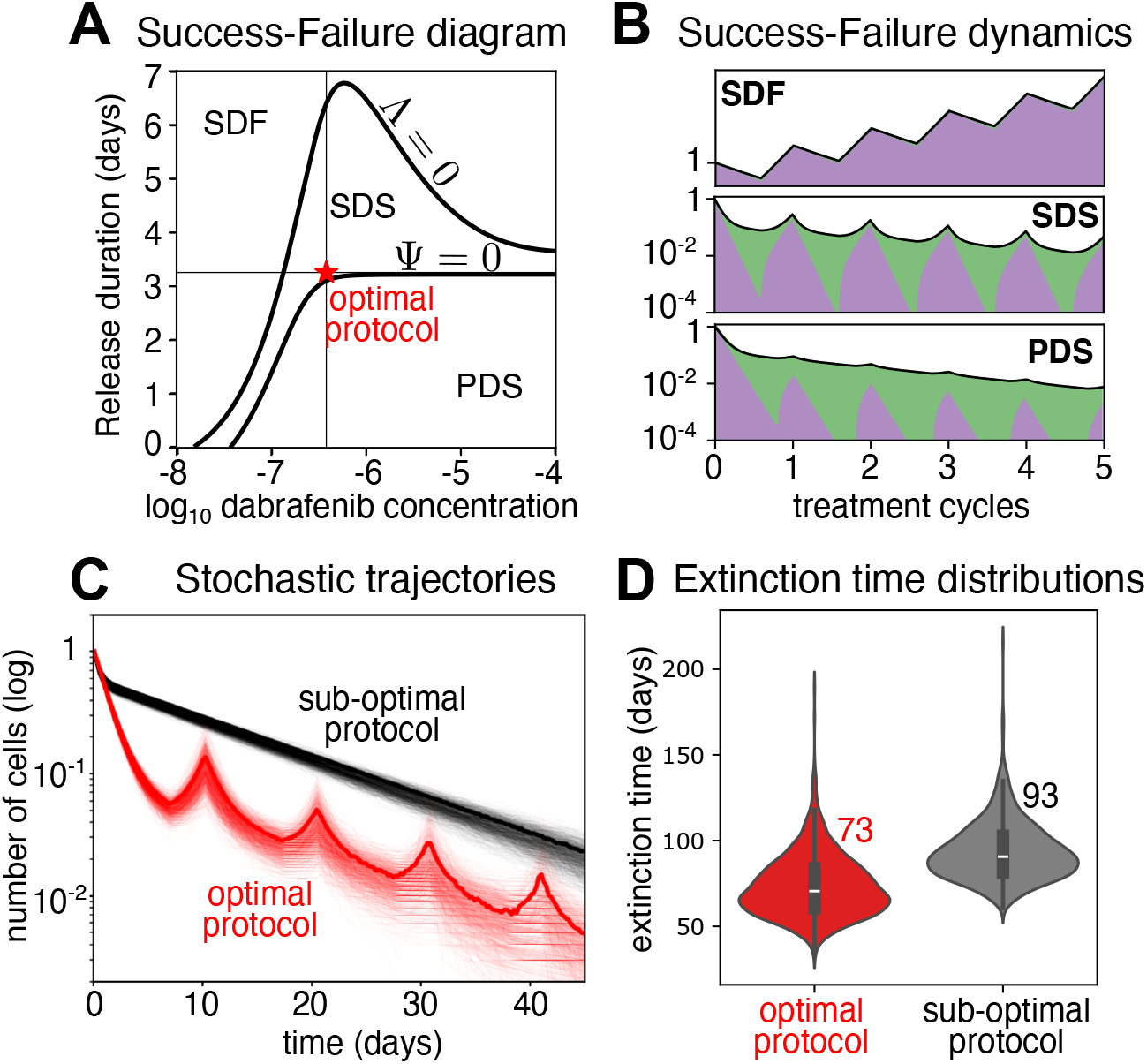
Conditions for the design of successful periodic protocols at constant treatment time. (A) Success-Failure diagram recapitulating the conditions for which specific choices of drug concentration and release duration, at fixed treatment duration, will bring the cancer population to extinction and with which subpopulation of cells (susceptible or persister cells). As depicted in Fig. 3 of the main text, phase boundary curves divide the plane in the same three distinct regions: “susceptible-driven failure (SDF)”, “susceptible-driven success (SDS) and “persister-driven success (PDS)” regions. The best treatment strategy minimizing the long-term fitness, depicted as a red star, is obtained at values (*τ*_*t*_ = 7 days, *τ*_*r*_ = 3.3 days, *c* = 3.8 *×* 10^−7^ *M* ). (B) Model’s predicted dynamics under protocols in each of the three regions, confirming the expected different behaviors. (C) Simulation of 1000 stochastic trajectories under the optimal protocol (red lines), and a sub-optimal protocol (black lines) in which drug concentration is 10x higher than clinical and release time is 0 (continuous treatment). Starting population size was set to 1000 cells. (D) Extinction time distributions computed from the simulated stochastic trajectories show that optimal protocol (in red) has a faster extinction dynamic compared to a continuous (zero release time) protocol in which 10x than clinical drug dose was delivered (in black). Average extinction times are approximately equal to 73 days, for the optimal protocol, and 93 days for the suboptimal protocol.

**Supplementary Fig. 12.**
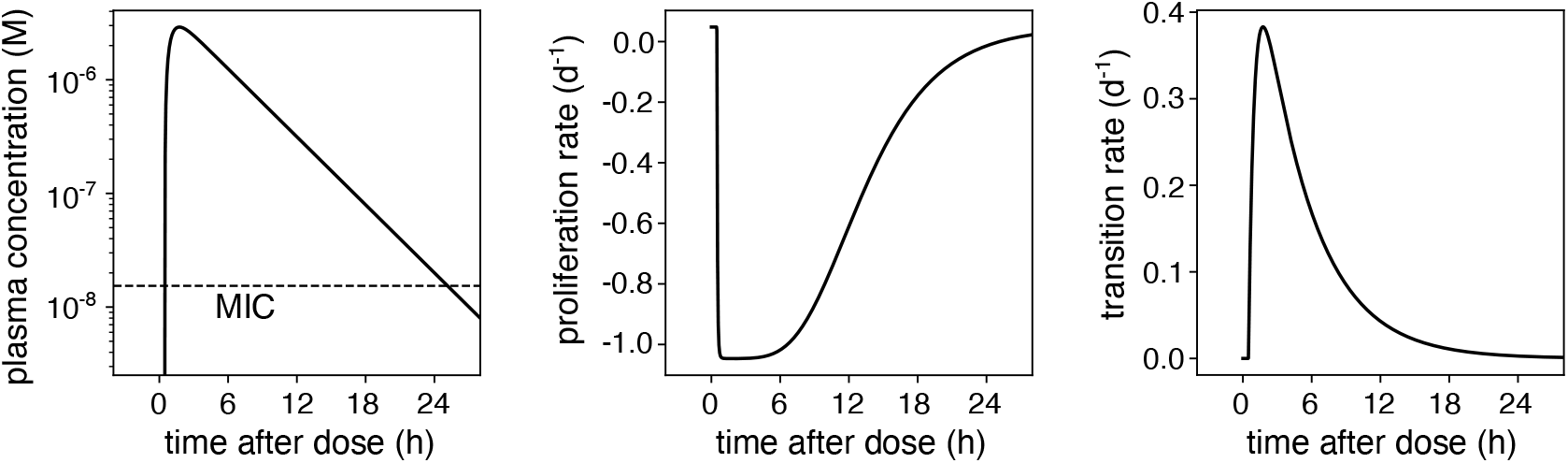
The dabrafenib dose profile felt by a patient induces a temporal dependence of both transition to persister and persisters’ death rates. (A) Dabrafenib plasma concentration temporal dependence inferred from patients’ drug pharmacokinetics data from ref. [18] of the main text. (B) Temporal dependence of persister cells’ proliferation (death) rate induced by the plasma concentration dose profile, as predicted by Eq. (4) of the main text. (C) Temporal dependence of transition to persistence rate, induced by the plasma concentration dose profile, as predicted by Eq. (3) of the main text. WiDr cl. B7 parameters.

